# Harmonization of the fastest and densest responses embodies the humanlike reaction time of mice

**DOI:** 10.1101/2024.04.17.590007

**Authors:** Chan Hee Kim

## Abstract

Reaction time (RT) is important in evaluating delayed latency in behavior. Unlike that of humans, the RT of animals, in which the stimulus–response relationship is not one-to-one due to repeated responses per trial, may exhibit two peaks of the fastest and densest responses in a distribution of responses. We determined whether the two peak latencies are aligned for a single RT by controlling stimulus duration. In delay conditioning with mice using sound cues of 10, 5, and 2s, the 2s group exhibited the strongest positive correlations between the two peaks, as well as responses’ number and accuracy rate, suggesting coupling of the fastest and densest responses, and a one-to-one relationship between stimulus and response, respectively. We propose the use of harmonization of the two peaks, elicited by stimuli that induce minimal responses, as a criterion for designing animal experiments to mimic humanlike RT.

## Introduction

Timing is critical for investigating behavioral response^1^. Reaction time (RT), also known as response time^2-5^, is an essential component in behavior analysis and serves as a marker reflecting each subject’s backgrounds^6^, sensory and cognitive processes^7-14^, and clinical perspective^15,16^. RT is also associated with neural correlation across multiple brain regions^17-19^. In human and animal studies, terms such as reaction speed^20,21^ and response latency^22,23^ are utilized interchangeably with RT. However, subtle methodological differences exist between the two due to their distinct behavioral characteristics, even when directly comparing the RTs between animals and humans^24-26^. Even if there are similarities between human and animal behaviors under different conditions, the results may be interpreted differently. For instance, animal behavioral responses have focused on whether animals sufficiently and successfully respond in experimental paradigms, and then the repeated responses have considered as part of animals’ nature. However, these repeated responses contain so many RTs, making it impossible to discern which ones are genuine. The shorter stimulus may reduce their unnecessary repeated responses. The current study proposes theoretical criteria for determining humanlike RT from animal data.

RT is typically evaluated within a specific time frame for each trial^27^, along with an estimate of its time point^28,29^. Researchers must provide clear instructions to human subjects before experiments to ensure RT sensitivity. Human subjects are occasionally asked to respond as fast as possible^30,31^. However, instructing animals to quickly respond may be impractical due to communication limitations. Even with clear instructions for animal subjects with a long training period in an experimental paradigm, animals may intentionally delay their responses. Furthermore, their behavior during a trial, which includes repeated actions such as licking, nose poking, and lever pressing, complicates the interpretation of an RT-like measure. In animal studies, the stimulus–response relationship per trial may not be one-to-one, as in human studies^32-34^. Whether the delayed latencies of repeated responses accurately reflect the animals’ decision-making points in the cognitive process is unclear. Averaged animals’ responses may contain errors in which the intention cannot be accurately determined. Direct comparison of an averaged latency in animals with RT in human studies would be difficult. Thus, the frequency of repeated responses, such as head entry (HE) into the food cup, is critical for assessing the animals’ behavioral performance^35-37^. During the training period, repeated HEs for repeated stimulations gradually become the densest point, forming a peak in the HE distribution that reflects the behavioral habituation and sensitization of subjects^38-43^. However, in addition to HE peak latency, the fastest HE (FHE) and its peak may exist within the HE distribution (Fig. 1 and S1). Fundamentally, the time points for RTs in animal data would not be a single latency, but rather a subset of both components. The coexistence of two distinct components may raise an issue that requires the identification of a true RT in the data. Considering one-to-one relationship like that in humans, the FHE peak of the first HE after stimulus onset would be the true RT. However, the HE peak would be more reliable as the true RT owing to its higher frequency. However, if the FHE and HE peaks coincide, it indicates a coupling of the fastest and densest responses. This is an ideal option that is closer to humans’ RT.

**Fig. 1.**
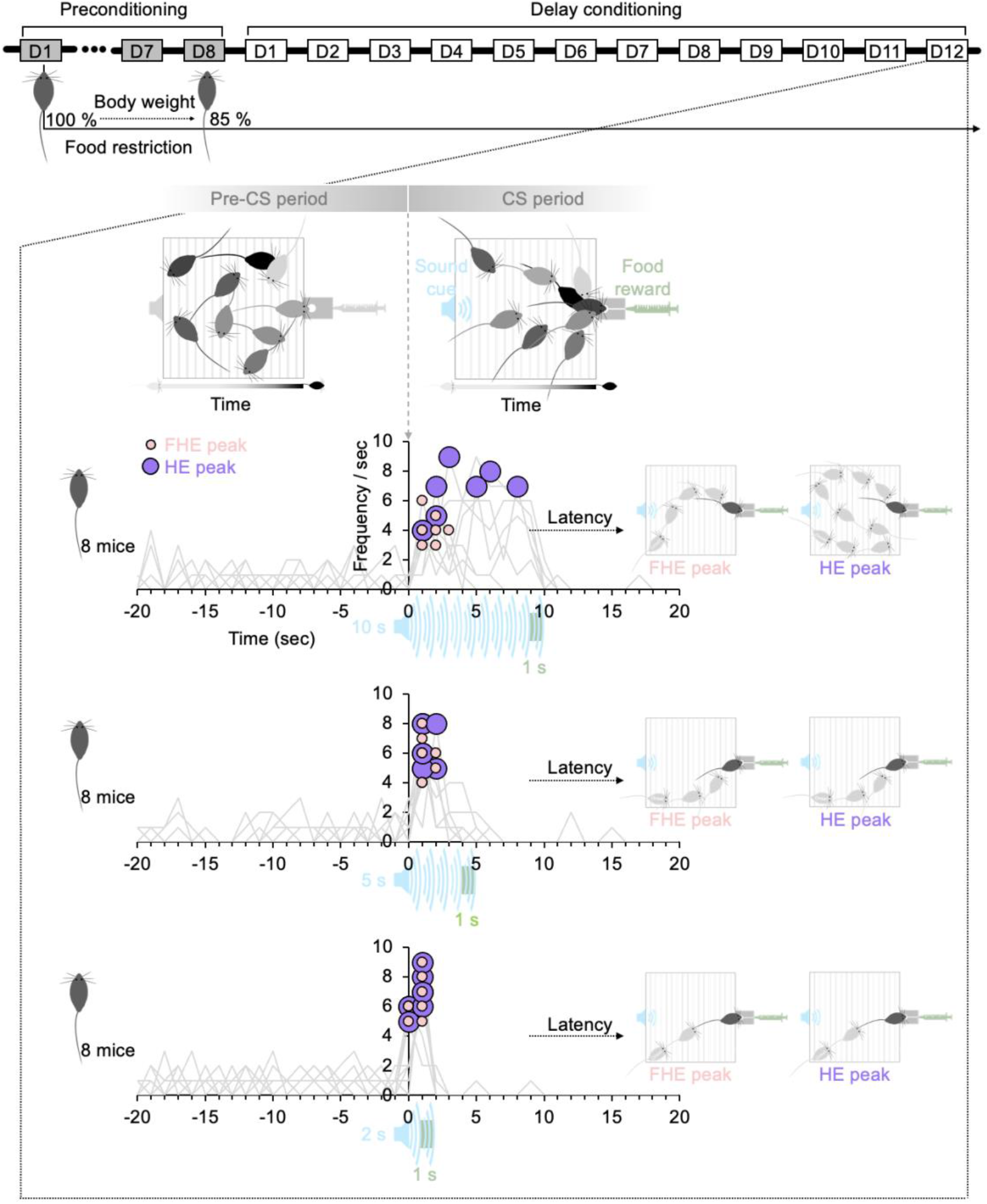
Experimental paradigm. The mice in all groups lost up to 85% of their body weight by restricting their food intake before delay conditioning. During the CS period, the duration of the sound cue followed by food reward varied among the 10-, 5-, and 2s groups. On the final training day (D12), the HE peak latency in the CS period after the onset of the sound cue was higher than that in the pre-CS period before the onset of the sound cue. The FHE peaks occurred before the HE peaks in the 10s group, whereas in the 5 and 2s groups, the two peaks were nearly synchronized. The latencies of the two peaks show how mice in each group respond differently to sound cues with varying durations. HE, head entry; FHE, fastest HE; D, training day; CS, conditioned stimulus

The temporal aspects^44-47^ of the experimental paradigm, such as the interstimulus interval, intertrial interval, interval between conditioned stimulus (CS) and unconditioned stimulus (US), and CS duration, have an individual and collective influence on the response. This study aimed to investigate the effect of CS duration on the determination of the CS–US interval in delay conditioning. In classical conditioning^48-50^, the time allowed for animals to respond is determined by the interval between the stimulus and the reward. The US of food following the CS onset of the stimulus cue guides the HEs. Some animals can delay their actions after deciding on long CS–US intervals. A short CS–US interval is preferable for eliciting more prompt responses from animals, whereas a long CS–US interval results in the safe and complete acquisition of more responses. A higher level of frequency on delayed latencies may indicate the animals’ behavior similar to food addiction^51^ as well as their deep awareness of task^52^. Latency and frequency are likely to exhibit greater distribution variance in long than in short CS duration, reflecting individual differences such as animals’ preferences, hesitation, and laziness, which deviate from humanlike RT. Indeed, many studies have explored the effect and significance of the CS–US interval (or CS duration) based on tests with various CS durations^35-37,45,46,53-58^. However, the specific durations of the stimulus vary depending on the apparatus and study paradigm, thus, an absolute value cannot be used to determine the appropriate duration for designing an experiment.

RT can be influenced by various experimental conditions. Our objective was to define the effects of a temporal factor in experimental design that could combine the two different latencies of FHE and HE peaks into a single cohesive unit of RT rather than treating them as separate entities. We thought that the factor was CS duration. To investigate this, we tested separate groups of C57BL/6 mice (eight male mice for each group) with sound cues of different CS durations (white noise of 10, 5, and 2 s) (Fig. 1). We hypothesized that the latency and frequency of the FHE and HE peaks would vary depending on the CS durations. If the FHE and HE peak latencies are synchronized, their frequencies would also align into a single peak because the relationship between latency and frequency can be interdependent. Moreover, we hypothesized that the optimal CS duration could be predicted based on the relationship between the two peaks. Thus, we recorded the HEs of freely moving mice in an operant chamber based on delay conditioning^59^, using white noise and condensed milk. To eliminate additional factors other than training food reward associations and to observe the mice’s natural behaviors throughout a sound cue, we did not include no-reward or punishment for missed responses in the experimental protocol. Rather than trace conditioning related to memory-based process^60^, delay conditioning was employed to elicit a response based on stimulus duration without an interval of silence.

In the data analysis, we conducted a detailed analysis of behavioral data at both the temporal and frequency levels to fully evaluate the impact of CS duration. Our analysis encompassed several steps. First, we assessed the difference and correlation between the FHE and HE peaks to better understand their relationship. Second, we separately investigated the group differences in the FHE and HE peaks to better understand their characteristics and implications. Third, we examined classical components such as HE frequency during the CS and pre-CS periods, as well as HE accuracy to confirm the effect of CS duration on subject performance. Furthermore, we considered how shorter compared with longer CS durations affected the mice’s ability to reach the food cup as well as how CS duration affected the HE frequency. Based on these findings, we determined how subjects’ performance was affected by the relationship between FHE and HE peaks. Finally, we examined the correlation between the number of HEs for each trial and accuracy across each group to determine whether the relationship between HE responses and correctness reflected humanlike behavior. Finally, we proposed a criterion for determining an appropriate CS duration to elicit a humanlike RT based on the alignment of the FHE and HE peaks.

## Results

### Difference between the FHE and HE peaks for each group

The nonparametric paired t-test results for all training days indicated that the 10s group mostly exhibited differences in latency and frequency between the FHE and HE peaks. The significance of these findings varied by training days (D). Specifically, for latency, the HE peaks outperformed the FHE peaks on D5, D7, D8, D9, and D11 (Fig. 2A and Table S1; Wilcoxon signed-ranks test, *P* < 0.05 in all cases). Similarly, for frequency, the HE peaks surpassed the FHE peaks on D5 and D7 through D12 (Fig. 2B and Table S2; Wilcoxon signed-ranks test, *P* < 0.05 in all cases). In 2s group, a significant difference was observed in D3. The 5s group was only significant in D11. Except for the aforementioned results, no significant differences were observed in latency or frequency between the FHE and HE peaks (P > 0.05 in all cases).

**Fig. 2.**
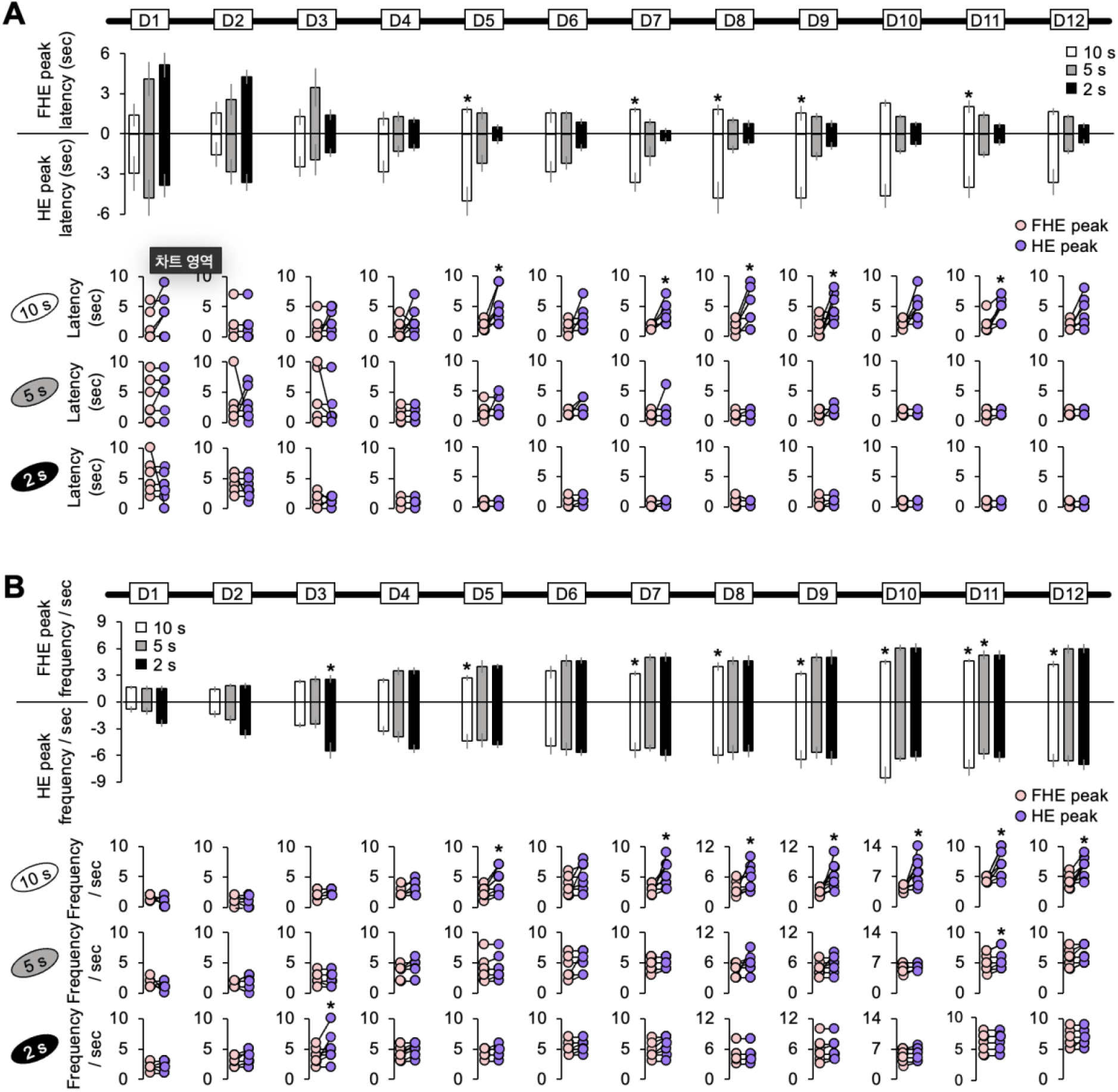
Difference between the FHE and HE peaks in latency and frequency. **(A)** Only the 10s group exhibited significant differences in latencies between the FHE and HE peaks (Wilcoxon signed-ranks test, *P* < 0.05). The latency for the HE peak was greater than that for the FHE peak. However, no significance was observed in the 5 and 2s groups (Wilcoxon signed-ranks test, *P* > 0.05). **(B)** The 10s group showed significant differences in the FHE and HE frequencies in D5 and D7–D12 (Wilcoxon signed-ranks test, *P* < 0.05). The frequency of the HE peak was higher than for the FHE peak. In 2s group, a significant difference was observed in D3. The 5s group was only significant in D11. See also Tables S1and S2 for the details of statistical analyses. *, *P* < 0.05. HE, head entry; FHE, fastest HE; D, training day

### Correlation between FHE and HE peaks for each group

We employed the Spearman test to determine the correlation between the FHE and HE peaks, based on the nonparametric paired t-test results. As expected, the FHE and HE peak latencies of the 2s group were strongly and positively correlated during the later training days (D4–D6 and D8–D12; Spearman test, *P* < 0.05 in all cases). Similarly, in the 5s group, positive correlations were observed on D1, D4, D8, D10, D11, and D12. For the 10s group, D1 and D2 were the only significant variables (Fig. 3A and Table S3). For the frequencies of the FHE and HE peaks, the 2s group exhibited positive correlations across all 12 days (Spearman test, *P* < 0.05 in all cases), except for D3 (Fig. 3B and Table S4). Significant positive correlations were also found in the 5s group on all days except for D1, D8, and D9. In the 10s group, the significant results were observed only in the early training period of D3 and D5, with a significant negative correlation on D1. The correlation results indicate that the FHE and HE peaks were similar in groups with short and shorter CS durations (2 and 5 s), but not in the group with long CS duration (10 s). In particular, the two peaks nearly overlapped in the 2s group.

**Fig. 3.**
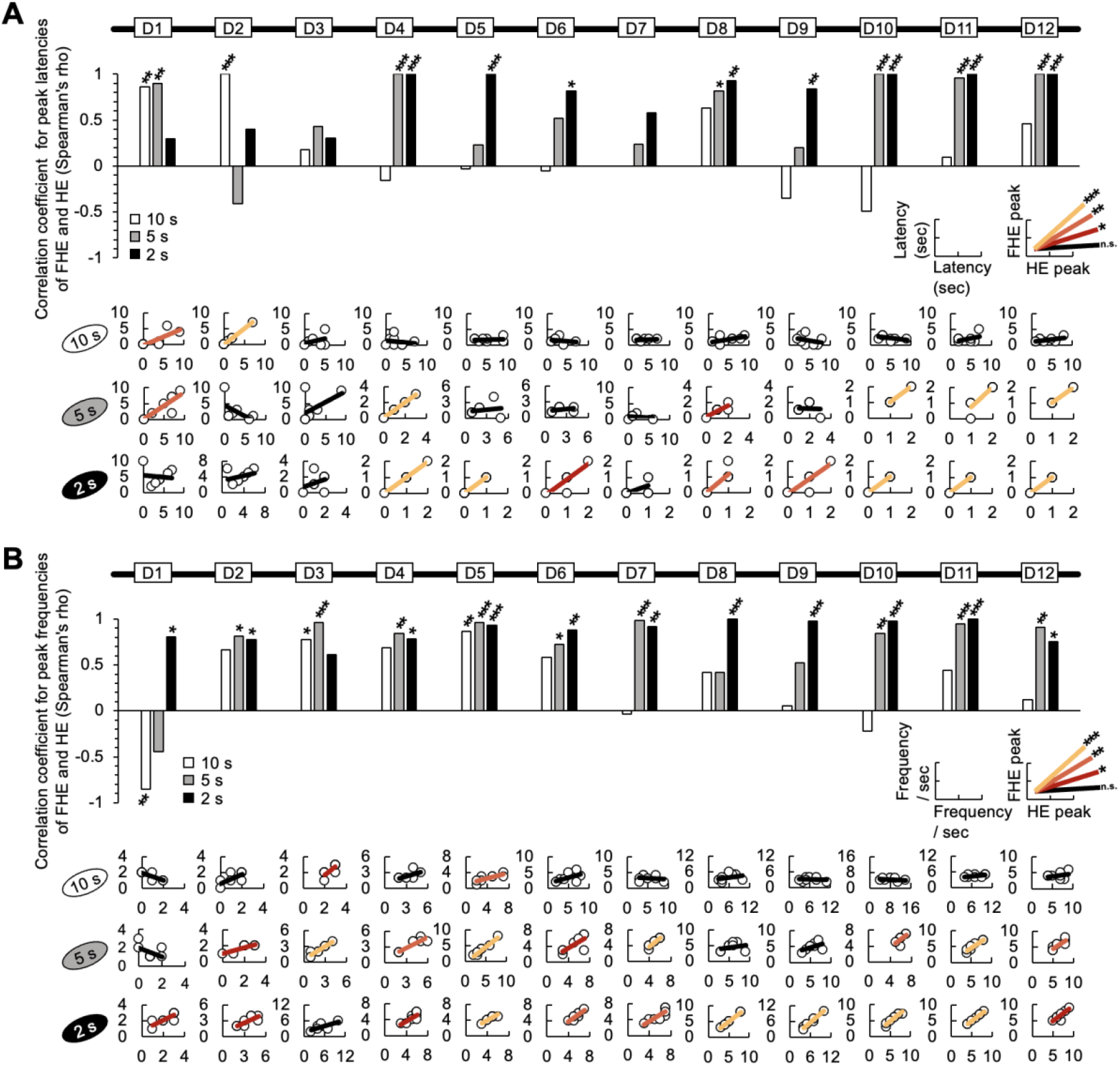
Correlation between the FHE and HE peaks in latency and frequency. **(A)** In the results of Spearman test, the latencies of the FHE and HE peaks were strongly and positively correlated in later training days for the 2s group (D4–D6 and D8–D12). Positive correlations were found in the later days of the 5s group (D1, D4, D8, D10, D11, and D12), but only on D1 and D2 for the 10s group. **(B)** Spearman test exhibited that the 2s group exhibited strong positive correlations in the frequencies of the FHE and HE peaks across all 12 days, except for D3. Furthermore, significant positive correlations were observed in the 5 and 10s groups. In the 5s group, positive correlations were found on all days except for D1, D8, and D9. However, in the 10s group, positive correlations were observed only during the early stages of training (D3 and D5). The nonparametric correlation was evaluated using the Spearman test. See also Tables S3and S4 for the details of statistical analyses. *, *P* < 0.05; **, *P* < 0.01; and ***, *P* < 0.001. HE, head entry; FHE, fastest HE; D, training day

### Group difference in FHE and HE peaks

In the results of one-way nonparametric analysis of variances (ANOVAs) with the group factor (10, 5, and 2 s) using the Kruskal–Wallis H test, the FHE peaks exerted a significant group effect across the training days for both latency (in cases of D1, D2, D5, D7, D10, D11, and D12) and frequency (in cases of D2, D3, D4, D5, D7 D9, D10, and D12) (Fig. 4A). Post hoc revealed significant differences mostly between the 10 and 2s groups or between the 10 and 5s groups for both latency and frequency except for D2 in frequency (Dunn’s Test, Bonferroni-corrected *P* < 0.05; Tables S5 and S6).

**Fig. 4.**
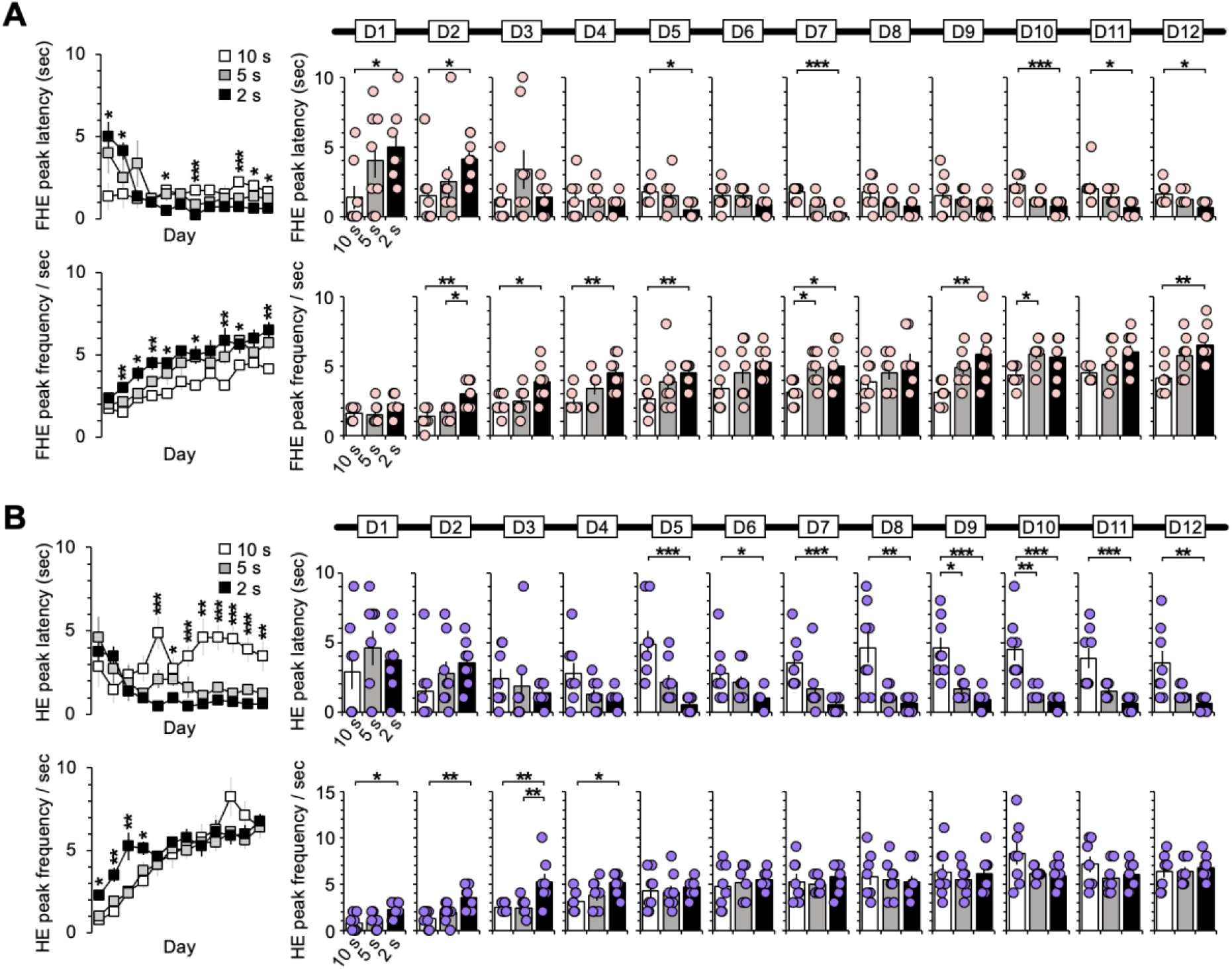
Difference between groups in the latency and frequency of the FHE and HE peaks. The one-way ANOVA results for the FHE and HE peak latency and frequency indicated a significant effect of group factor (10, 5, and 2 s). **(A)** The FHE peak exhibited a significant group effect across training days for both latency (D1, D2, D5, D7, D10, D11, and D12) and frequency (D2, D3, D4, D5, D7, D9, D10, and D12) (Kruskal–Wallis H test, *P* < 0.05). Post hoc analyses revealed significant differences between the 10 and 2s groups or between the 10 and 5s groups for both latency and frequency (Dunn’s Test, *P* < 0.05, Bonferroni-corrected). **(B)** The one-way ANOVA results for the HE peak indicated a significant group effect for both latency (D5–D12) and frequency (D1–D4) (Kruskal–Wallis H test, *P* < 0.05). In the post hoc analysis, the latency of the 10s group significantly differed from the other groups starting with D5 (Dunn’s Test, *P* < 0.05, Bonferroni-corrected). In the post hoc analysis for frequency, the 2s group differed the most from the other groups from D1 to D4 (Dunn’s Test, *P* < 0.05, Bonferroni-corrected). See also Tables S5 to S8 for the details of statistical analyses. *, *P* < 0.05; **, *P* < 0.01; and ***, *P* < 0.001. Error bars denote 95% confidence intervals. HE, head entry; FHE, fastest HE; D, training day

Meanwhile, one-way ANOVA results for the HE peaks revealed a pattern in which the significant group effect for latency on D5–D12 appeared to mirror the significance observed in frequency on D1–D4 (Fig. 4B). Post-hoc analysis for latency revealed significant differences between the 10s group and other groups during training periods from D5 to D12 (Dunn’s Test, Bonferroni-corrected *P* < 0.05; Table S7). In the post-hoc analysis for frequency, the group differences were significantly higher in the 2s group than in other groups during the early training periods of D1 to D4 (Dunn’s Test, Bonferroni-corrected *P* < 0.05; Table S8).

Based on the results of the FHE and HE peaks, the group differences in the FHE peaks were sporadic and irregular across the training days whereas those in the HE peaks followed a more consistent pattern. We concluded that the results in the HE peak reflected stimulus adaptation after D1 to D4, as differences in the HE peak latency between the longest and shortest CS groups persisted from D5 to D12. Our results indicate that the HE peaks explain the effect of CS duration better than the FHE peaks.

### HE accuracy

The HE accuracy was considered to be correct if the mice’s heads touched the food cup within the CS duration of 10, 5, and 2 s. The results of the one-way ANOVAs using Kruskal– Wallis H test indicated significant group differences in HE accuracy on D7, D8, D9, and D11 (Fig. 5A). Post hoc analyses revealed only differences between the 10 and 2s groups (Dunn’s Test, Bonferroni-corrected *P* < 0.05; Table S9). However, no significant difference was discovered on the last day of D12. Overall, the subjects achieved high accuracy rates of more than 80%, regardless of the CS duration.

**Fig. 5.**
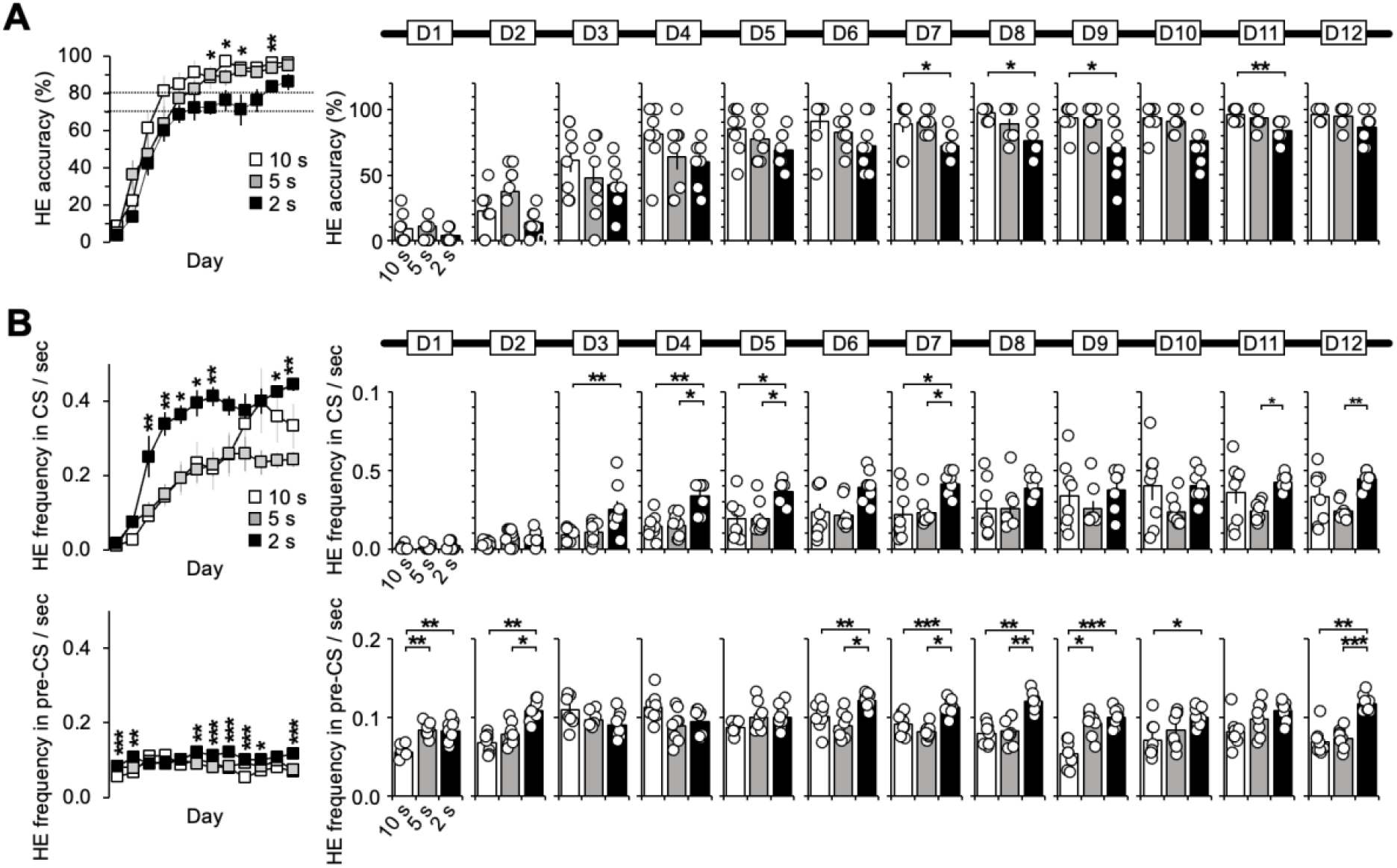
Basic properties of HE. (**A)** The one-way ANOVA results indicated that the group effect for HE accuracy was only present on D7, D8, D9, and D11 (Kruskal–Wallis H test, *P* < 0.05). However, all subjects eventually performed well on the tasks, with accuracy above the mean of 80%, regardless of the CS duration. **(B)** One-way ANOVA revealed a significant group effect for HE frequency during the CS period (D3–D5, D7, D11, and D12) (Kruskal– Wallis H test, *P* < 0.05). Post hoc analysis revealed that the 2s group had a significantly higher frequency than the 5 and 10s groups (Dunn’s Test, *P* < 0.05, Bonferroni-corrected). During the pre-CS period, the group effect was significant on 8 days except for D3, D4, D5, and D11 (Kruskal–Wallis H test, *P* < 0.05). Post hoc analysis revealed that the 2s group had higher frequencies than the 10s group. D1 and D9 showed difference between the 10 and 5s groups, while D2, D6, D7, D8, and D12 revealed differences between the 5 and 2s groups (Dunn’s Test, *P* < 0.05, Bonferroni-corrected). See also Tables S9 to S11 for the details of statistical analyses. *, *P* < 0.05; **, *P* < 0.01; and ***, *P* < 0.001. Error bars denote 95% confidence intervals. HE, head entry; FHE, fastest HE; D, training day; CS, conditioned stimulus

### HE frequency

HE frequency was calculated for both the CS and pre-CS periods, with the number of HEs normalized per 1 s for the three groups. During the CS period, significant group differences were observed on D3–D7, D11, and D12 (Fig. 5B and Table S10). Post hoc analysis revealed that the 2s group exhibited significantly higher frequencies than the 5 and 10s groups in D3, D4, D5, D7, D11, and D12 (Dunn’s Test, Bonferroni-corrected *P* < 0.05). For the pre-CS period, one-way ANOVAs results indicated significant group effects on 8 days except for D3, D4, D5, and D11 (Fig. 5B and Table S11). In post hoc, the 2s group exhibited higher frequencies than the 10 and 5s groups. The 10 and 2s groups showed differences on the days except for D3, D4, D5, and D11, the 10 and 5s groups showed differences on D1 and D9, and the 5 and 2s groups differed on D2, D6, D7, D8, and D12 (Dunn’s Test, Bonferroni-corrected *P* < 0.05). These findings show how CS duration affects the pre-CS resting state. Different CS durations most likely resulted in frequent or infrequent HEs from the subjects.

Additional analysis was conducted on baseline-corrected values of HE frequency in the CS and HE peak. The mean values of HE frequency and HE peak in the pre-CS period were used as the baselines, and each baseline effect was subtracted from HE frequency in the CS and HE peak. However, there was no prominent difference from the uncorrected results (Fig. S2 and Tables S12, S13, and S14). We concluded that the responses in the off-stimulus of pre-CS did not directly impact the HE peak and frequency in the on-stimulus of CS.

### Number of HE and correlation with accuracy

For each group, we counted the number of HEs in the CS period without normalizing per 1 s (as we did with HE frequency). As expected, longer CS durations elicited a higher response rate from subjects (Fig. S3A). In the 10s group, the HE occurrences increased to approximately three after D8. Contrarily, in the 2s group, the HE occurrences remained below 1, whereas in the 5s group, HE slightly surpassed 1. On D12, the mean and standard deviation values were 3.325 ± 1.676 in the 10s group, 1.123 ± 0.285 in the 5s group, and 0.888 ± 0.099 in the 2s group. Significant group differences were observed between 10 and 2s in D4 and D8–D12 (Dunn’s Test, Bonferroni-corrected *P* < 0.05; Table S15).

Next, we explored the relationship between the accuracy rate and the number of HEs per trial, which was normalized to a percentage value for an easier comparison with the accuracy rate. Each HE occurrence was scored as ten. The results were highly significant across all training days, except for D7 in the 2s group (Spearman test, *P* < 0.05; Figs. 6A and S3B). The 2s group exhibited a linear trend in which the accuracy rate improved with an increase in the number of HEs across all training days (Fig. 6B and Table S16), with the fewest HEs occurring less than once per trial (Fig. 6C). However, the correlation results for the 5 and 10s groups varied across training days and became less significant near the final day. The lack of significance in the correlation results for the 10 and 5s groups could be attributed to the accuracy rate ceiling effect, or to an excess of HEs relative to the number of trials.

**Fig. 6.**
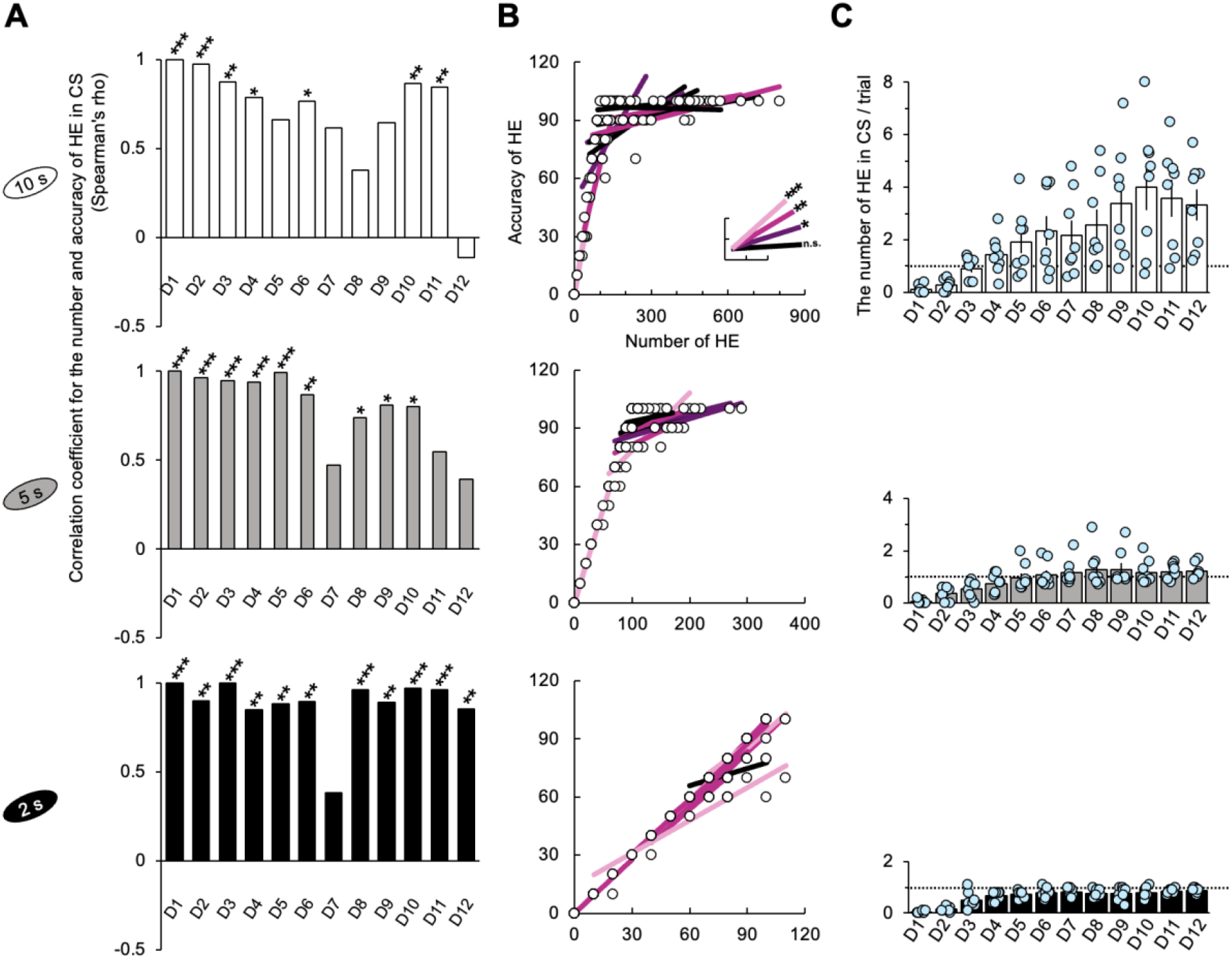
Number of HE and correlation with accuracy. **(A)** Significant positive correlations were observed between the number of HEs and accuracy across all groups (Spearman test, *P* < 0.05). For the 2s group, these significances were consistent across all the training days, except for D7. The nonparametric correlation was assessed using the Spearman’s test. **(B)** The graphs included the lines of best fit and data points corresponding to the correlation coefficients in Fig.6A and show the trend in the data for each day (Fig. S3B). The scale of the x-axis was determined by the maximum value observed across all the training days. In the case of the 2s group, many data points overlap, resulting in a small representation of the 96 points collected from 8 subjects over 12 days. **(C)** The 10s group exhibited the highest number of HEs. In the 2s group, the HE occurrences were below 1, while in the 5s group, HE slightly exceeded 1. The dotted line represents the number of HE that corresponds to the ono-to-one matched relationship between stimulus and response. See also Tables S15 and S16 for the details of statistical analyses. *, *P* < 0.05; **, *P* < 0.01; and ***, *P* < 0.001. Error bars denote 95% confidence intervals. HE, head entry; FHE, fastest HE; D, training day; CS, conditioned stimulus

### Repetition of HE

In the 2s group, the one-to-one matched relation may have occurred due to the mice’s restrained impulsivity for anticipated food reward, or it may simply reflect their natural tendency to momentarily pause after reaching the food cup. To explore these possibilities, we sorted the HEs for each trial chronologically and calculated the time intervals between them: from the CS onset to the first HE, from the first to the second HE, and between the subsequent HEs. We counted the number of mice involved in each HE order as well as the frequency with which responses occurred. If a mouse responded at least once in ten trials, it was counted as 1 mouse in the order of that HE. For each order, the number of HEs was normalized to the percentile, with 80 responses to 80 trials equaling 100%. Because of incomplete successive observations of responses beyond the second HE for all subjects, statistical analysis could not be performed in this study.

The maximum number of HE was ten times across all trials and training days (Figs. 7B and S4). In the 10s group, one mouse responded up to ten times on D9 and D10 (10^th^ HE in Fig. S4B). The seventh HE in the 5s group (7^th^ HE in Fig. S4B) and the third HE in the 2s group were the highest recorded (3^rh^ HE in Fig. 7B). The number of subjects who produced their first HE increased by D12 across all groups. In addition, up to eight mice of all subject number in both the 10 and 5s groups produced the second HE by D12, indicating an association between the first and second HEs. However, this pattern did not appear in the 2s group (Fig. 7A). The second HE of the 2s group from D3 to D7, which had a similar number of participating subjects to the 5s group, was faster than that of the 5s group, with intervals mostly ranging from 0 and 1 s. However, the number of subjects did not increase as they approached D12, unlike the 5s group. Despite responding similarly during the 2s CS period, the mice in this group exhibited fewer repeated HEs than the other groups. These findings indicate that the mice in the 2s group learned the stimulus–response relation well. Contrarily, the mice in the 10 and 5s groups demonstrated more variable responses across training days, as evidenced by fluctuations in the correlation coefficients between the number of HEs and accuracy rate.

**Fig. 7.**
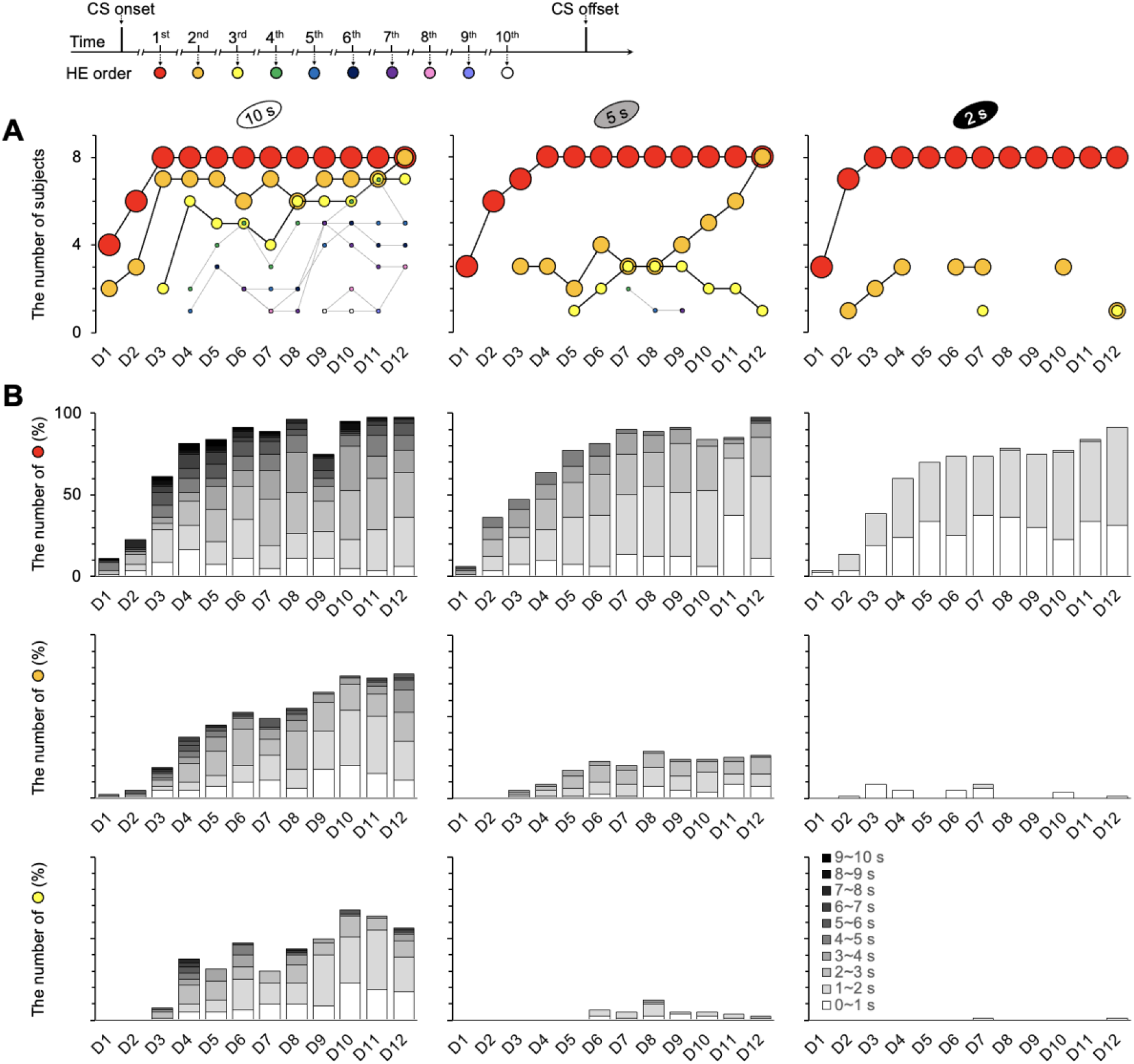
Number of repeating HEs and participating subjects per trial. The repetitions of HEs were observed up to ten times across all trials and training days. **(A)** In the 10s group, the responses were recorded until the tenth HE (Fig. S4B). Meanwhile, the maximum observed in the 5s group was the seventh HE (Fig. S4B), while in the 2s group, it was the third HE. The pattern of the first HE was consistent across all groups; however, differences were observed in the second and third HEs. Specifically, the second HE increased toward eight mice on D12 in both the 10 and 5s groups but not for the 2s group. The second HE of the 2s group from D3 to D7, which had the same number of participating subjects as the 5s group, was faster than the 5s group, with intervals ranging from 0 and 1 s in Fig. 7B. However, it did not increase as the 5s group toward D12. **(B)** The latencies between HEs varied with the CS durations. The interval latency was the longest in the 10s group and the shortest in the 2s group. See Fig. S4 for the details of the results from the fourth through the tenth HEs for the 10 and 5s groups. HE, head entry; D, training day; CS, conditioned stimulus

## Discussion

In the current data, between the FHE and HE peaks, which could be considered as RT, it was difficult to determine which one is closer to RT. Positive correlation results between FHE and HE peaks provided a solution to the dilemma of deciding between the two peaks. As the CS duration decreased, the FHE and HE peak latencies became more aligned, which resulted in fewer HE occurrences and increased monotony, similar to human responses. The basic properties of HE accuracy and frequency showed that the mice could reach and respond to the food cup despite varying CS durations. The HE accuracy scores exceeded 80% for all the 10-, 5-, and 2-s groups. The HE frequency in the short and shorter CS durations (5 and 2 s) was the same as in the long CS duration (10 s). The group differences in pre-CS didn’t affect the results in the CS period or HE peaks.

In addition to accuracy, the latency of RT accesses subjects’ task perception^2^. In human studies^15,17^, accuracy and RT are extracted and compared using the same data point. However, in animal studies, subjects may respond repeatedly, resulting in multiple RTs across multiple data points, forming a single accuracy rate. RT may be a more sensitive indicator than accuracy owing to their different mechanisms^61^, which can greatly fluctuate depending on the researcher’s instructions. Subjects can be asked to slowly or quickly respond with little effect on accuracy scores. In our study, all three groups responded well to each stimulus (Fig. 5A), but the FHE and HE peak latencies differed (Fig. 4), indicating that different CS durations reflected the researchers’ instructions on response timing. Longer CS durations gave subjects more freedom in response timing, whereas shorter durations may have caused anxiety about responding within a limited time frame. Thus, the shorter CS duration of 2 s in our study represented good instruction for more sensitive RT, in which the FHE and HE peaks were nearly coincident in terms of latency and frequency and accuracy rate and HE number were positively correlated over the training periods.

The FHE and HE peaks exhibited distinct patterns between the groups. The HE peak clearly demonstrated the effect of different CS durations on latency and frequency across early to late training days, indicating a strong relationship with animals’ habituated behaviors for the stimulus^38-42^. Contrarily, the FHE peak showed rarely significant group differences over several days. The frequencies serve as reliability indicators, reflecting the overlapping responses of individual subjects at specific time points^35-37^. The nonsignificant group difference in the HE peak frequency after D5 (Fig. 4B) corresponded to the saturation of the HE frequency (Fig. 5B). The results of the HE peak indicate that all subjects in the three groups adapted to the CS duration and performed well within each CS period. However, the FHE peak results indicate that the response of the longer CS group was dispersed even within the subject’s data because of the low density of responses during the extended CS period compared to the shorter CS groups. The group differences in FHE and HE exhibited dissimilarity in latency and frequency. Based on our findings, we proposed that the 2s CS duration, which displays synchronized FHE and HE peaks in the current experimental paradigm, is the best CS duration for the attainment of unbiased results in mice behavior.

The 2s group had the highest frequency during the pre-CS period, indicating that they checked the food cup more frequently during shorter CS periods. In addition, a linear relationship was observed between the HE frequency and CS duration during the pre-CS period. However, it is worth nothing that behavior during the pre-CS period did not influence behavior during the on-stimulus period, as evidenced by the baseline-corrected results for the HE peak frequency and latency during CS. Despite having the highest number of HE occurrences during the CS period, the 10s group had no significant differences in HE frequency compared to the 5 or 2s groups. A linear relationship between CS duration and response^45,58^ was only observed between the 5 and the 2s groups. Contrary to the linear trend in the pre-CS period, the non-linear trend in the CS period suggests that CS duration plays a different role during on- and off-stimulus conditions.

The HE frequency during the CS period in the 5s group was significantly lower than that in the 2s group. This indicates that the response pattern of the 5 and 2s groups after the onset of the sound cue would most likely be similar. Interestingly, the latency and frequency of FHE and HE did not significantly differ between 5 and 2s groups, except for the HE peak frequency in D3 (Fig. 4). The 5 and 2s groups showed positive correlations between the FHE and HE peaks. However, the 2s group, which exhibited a stronger correlation between the FHE and HE peaks, also showed a correlation between the number of HE and accuracy over the training days. Meanwhile, the correlation results of the 5s group indicated that the mice’s late-day behaviors were subtle and arbitrary, similar to those of the 10s group. Furthermore, the 2s group responded less than 1 per trial, whereas the 10 and 5s groups responded with more than 1 HE. These findings suggest that the 5 and 2s groups performed similarly; however, the mice in the 2s group learned the one-to-one matched relationship between stimulus and response more effectively. Moreover, the results of the second HE indicated that the mice that produced the first HE repeated it at least once for any trial, but this was observed only in the 2s group out of all the groups (Fig. 7). Our findings suggest that the mice in the 2s group chose to stop responding during the remaining time until the US was presented after the first HE in the CS period. Contrarily, the 5s group exhibited a steep linear increase in the number of subjects involved in the second HE beginning with D8 and eventually reaching all subjects with D12, as did the 10s group. The synchrony between the FHE and HE peaks, as well as between the number of HEs and accuracy in the 2s group, was caused by the absence of the second HE. In our data, the prediction for the US encompassed not only enhancement of response^62-64^ but also both inhibition and promotion of response based on CS duration. We propose that the second HE, which is influenced by CS duration, can be interpreted as an indicator of impulsivity toward the US in our data.

The mice’s RT was comparable to that of humans based on the relationship between the FHE and HE peaks. In our paradigm, the shorter 2s CS yielded a one-to-one matched relationship with response frequencies of less than 0.5 Hz. However, if the data involves licking behavior with higher frequency responses in a head-fixed model, the relationship may change, despite the presence of synchronization between the FHE and HE peaks. The one-to-one matched response would be tailored to our paradigm. Nevertheless, our findings indicated how the variety of personalities among animals can be managed in terms of CS duration. Setting a time limit for responses can promote genuine participation without hesitation, thereby improving the validity and reliability of the behavioral data. Finally, we propose a novel model with quantifiable criteria for the evaluation of the stimulus duration in RT-related experimental paradigms regardless of apparatus and specific characteristics of the animals.

This model is based on the harmony the between FHE and HE peaks as well as the relationship between the number of HEs and accuracy (Fig. 8). If animals are capable, the shorter CS durations, which can barely elicit one response per trial, may be preferable for eliciting humanlike responses with no repetition.

**Fig. 8.**
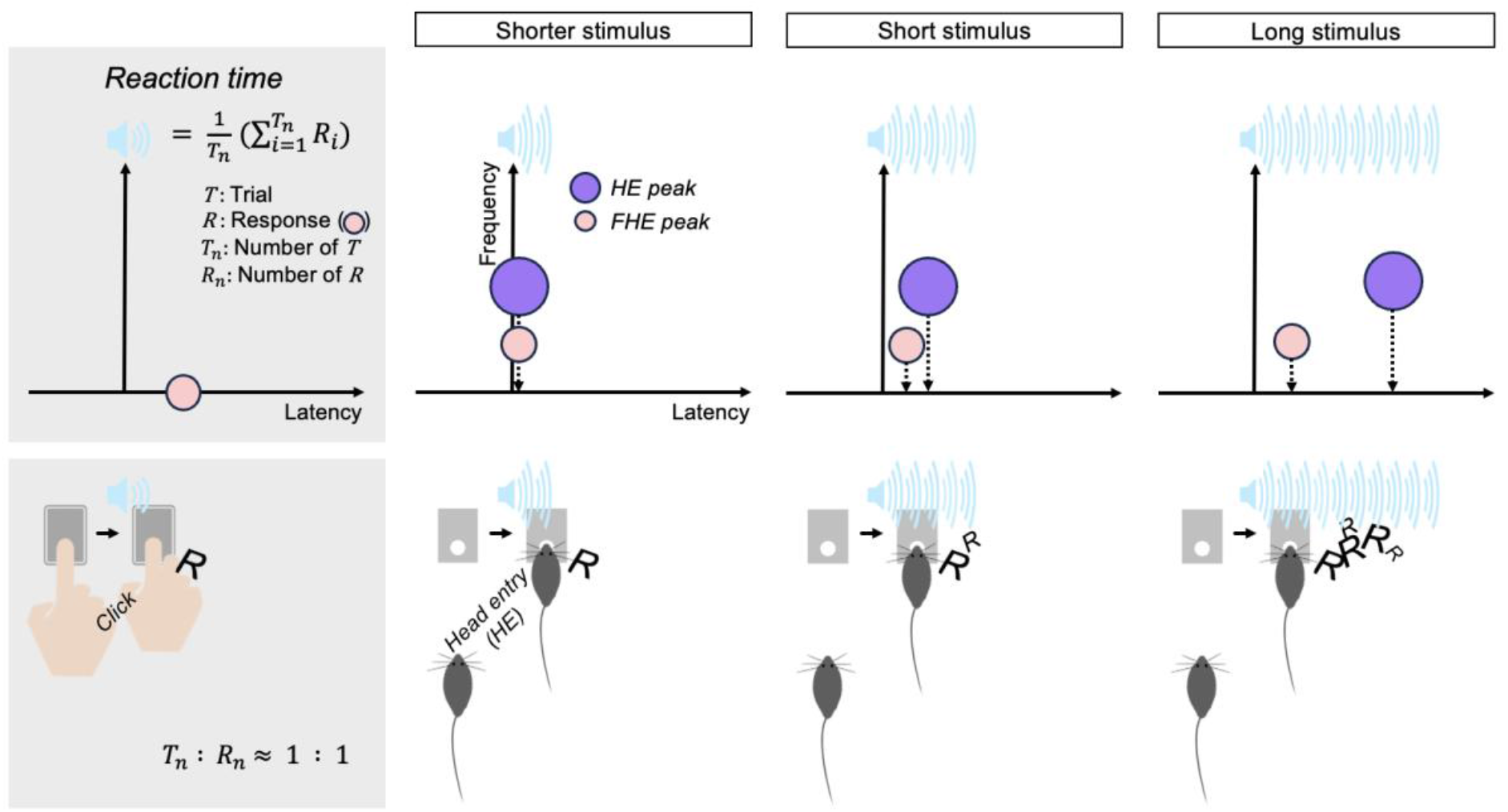
Reaction time model for mice. The harmonization of the HE and FHE peaks evoked by the shorter stimulus is associated with the humanlike response of mice. The number of responses to the shorter stimulus per trial is approximately 1, whereas the responses to the short and long stimuli are over 1, unlike humans. As for the number of responses, the computation of humanlike averaged RTs can be adapted only for the shorter stimulus. HE, head entry; FHE, fastest HE

## Methods

### Subjects

A total of 24 naïve male C57BL/6 mice, aged 7 weeks old, were provided by the Experimental Animal Center at Seoul National Dental University. Eight mice were assigned to each group. Before the delay conditioning, each mouse was housed in its own cage and subjected to a food restriction schedule to maintain their weight at 85% of their free-feeding weight throughout the study. They had unlimited access to water. The mice were kept in cages in a room with a light–dark cycle of 12 h each (8 am–8 pm). Training began when the mice reached 8 weeks old and ended when they were 10 weeks old. The care and use of every animal were approved by the Seoul National University Institutional Animal Care and Use Committee (SNU IACUC).

### Apparatus

The experiments were conducted using operant chambers (interior dimensions: 21.6 × 17.8 × 12.7 cm; model EBV-307W-CT, Med Associates), each equipped with a stainless steel grid floor consisting of 24 rods with a diameter of 0.32 cm (17.8 × 15.2 × 5.7 cm, model ENV-307W-GF) and enclosed within MDF sound-attenuating cubicles (model ENV-022MD). A house light (28 V DC, 1000 mA, model ENV-315W) was turned on from the start to the end of each experiment. A sound cue greater than 75 dB^65^ was produced by a white noise amplifier (10–25,000 Hz, model ENV-325SW) and delivered through a cage speaker (model ENV-324W). The noise produced by fans (model ENV-025F) in each cubicle was approximately 65 dB. Food reward, which consisted of condensed milk, was delivered into a single 0.5-cc stainless steel cup through a liquid pipe (model ENV-303LPHD) connected to a syringe pump (model PHM-100A-EURO). The HEs were counted whenever mice broke the beam of the detector (model ENV-303HDW) inside the liquid pipe. The experiments were conducted using the MED-PC V software (model SOF-735).

### Stimulus

The experiment included two stimuli: CS and US. The CS consisted of white noise cues with varying durations: 10, 5, and 2s (Fig. 1 and S5B). The selection of a CS duration of 10 s was influenced by literatures^36,45,66,67^ on CS duration. The 5 and 2s durations were determined using the HE and FHE peak latencies observed in our preliminary tests with a 10s CS duration. The FHE and HE peaks patterns were also observed in the current data (Fig. S5C). To ensure that the CS durations were within the ranges where mice could respond effectively, we examined the responses to the 10s CS. We selected the 2s duration based on the peak latency of FHE and the 5s duration based on the peak latency of HE observed during the late training sessions. The US consisted of condensed milk. In accordance with the principles of delay conditioning, the offset timings of the US matched those of the CSs (Fig. 1). Therefore, each US lasted 1 s and occurred between 9 and 10 s after the onset of the 10s CS, 4–5 s after the onset of the 5s CS, and 1–2 s after the onset of the 2s CS.

### Procedure

Before the main experiment’s delay conditioning phase, the preconditioning phase spanned 8 days, during which the mice were subjected to food restriction until their weight reached 85% of their baseline. They were also acclimated to handling and the US. On the last day of the preconditioning phase, the mice had a session in the chamber without the CS but with the US. After preconditioning, the delay conditioning experiments were repeated daily for a total of 12 days (Fig. 1). Each day, ten trials were conducted during the experiment. The Intertrial intervals varied and were randomly set to 90, 180, or 280, as illustrated in Fig. S5A. The total experimental time per subject was about 1 h per day, including the preparation time. During the experiments, pumps were used to deliver condensed milk from syringes to food cups, with the pumps turned on during the US period. Furthermore, the mice were free to move around the chamber. In this study, the behavior directed toward the food cup to consume the reward food was referred to as “HE.” The timing of HEs to the food cup was recorded with a temporal resolution of 10 ms.

### Analysis

The data was analyzed in three groups of 24 mice, each with duration of 10, 5, and 2s. For the HE frequency analysis, the HEs were extracted from the CS and pre-CS time windows for ten trials, and the number of HEs was normalized per s. We carefully selected the time window of pre-CS period as an off-stimulus baseline to compare to the CS of the on-stimulus condition. Fig. S6B shows the details of pre-CS segmentation. In the analysis of HE accuracy, the presence or absence of HE during each CS duration per trial was calculated and converted to percentages (where one correct trial equates to 10%). The number of HE was calculated using the total number of HEs during each CS duration. The FHEs were extracted from the first HE in each trial (Fig. S1A). The FHE and HE peaks were calculated by extracting maximum values from a matrix in which the numbers of HEs occurring within a 1-s epoch were recorded as frequency levels (Figs. S1B and S6). The maximum frequency value for each peak was 10, the trial number per day. The latency of the peaks corresponded to the time of maximum frequency. Accordingly, the latencies of the peaks did not include the incorrect HEs, as RT is typically based on correct answers^68,69^. The time windows for the estimation of the FHE and HE peaks were each set from 0 to 10 s after the onset of CS in all groups (Fig. S7). The baseline-corrected HE frequency and peak were calculated by subtracting the mean HE in the pre-CS period from the HE in the CS (Figs. S2 and S8).

### Statistics

The statistical analysis showed that all data did not follow a normal distribution. Thus, nonparametric analyses were employed for all statistical procedures. Nonparametric paired t-tests between the FHE and HE peaks were conducted using the Wilcoxon signed-ranks test. The nonparametric correlations between each pair of the FHE and HE peaks as well as the number of HEs and accuracy were determined using the Spearman test. All three groups had a correlation coefficient value of 1, suggesting that the FHE and HE peaks coincided (Table S4). Notably, the conversion of a decimal to an integer during the extraction of the two peaks from HEs exerted a significant effect on the highest correlation coefficient values (Fig. S6). Nonparametric ANOVAs were conducted using the Kruskal–Wallis H test. In addition, post hoc analyses for ANOVAs were conducted using the Dunn’s Test. Type I error caused by multiple comparisons between the three groups in the Dunn’s Test was adjusted using the Bonferroni test. MATLAB 9.12.0.2039608 (Math Works Inc., Natick, MA, USA) and SPSS 25.0 software (IBM, Armonk, NY, United States) were used to conducted all analyses.

## Data availability

The datasets used and/or analyzed during the current study are included in the article and supplementary information files, and are available from the corresponding author on reasonable request.

## Acknowledgement

This research was supported by Seoul National University Research Grant in 2021 and the Basic Science Research Program through the National Research Foundation of Korea (NRF) funded by the Ministry of Education (2022R1I1A1A01073800).

## References

1 Killeen, P. R. & Fetterman, J. G. A behavioral theory of timing. Psychol Rev 95, 274–295 (1988). 10.1037/0033-295x.95.2.274

2 Schouten, J. F. & Bekker, J. A. Reaction time and accuracy. Acta Psychol (Amst) 27, 143–153 (1967). 10.1016/0001-6918(67)90054-6

3 Luce, R. D. Response times: Their role in inferring elementary mental organization. (Oxford University Press, 1986).

4 Baayen, R. H. & Milin, P. Analyzing reaction times. International journal of psychological research 3, 12–28 (2010).

5 Welford, W., Brebner, J. M. & Kirby, N. Reaction times. (Stanford University, 1980).

6 Welford, A. T. Reaction time, speed of performance, and age. Ann N Y Acad Sci 515, 1–17 (1988). 10.1111/j.1749-6632.1988.tb32958.x

7 Wendt, D., Brand, T. & Kollmeier, B. An eye-tracking paradigm for analyzing the processing time of sentences with different linguistic complexities. PLoS One 9, e100186 (2014). 10.1371/journal.pone.0100186

8 Krajbich, I., Bartling, B., Hare, T. & Fehr, E. Rethinking fast and slow based on a critique of reaction-time reverse inference. Nat Commun 6, 7455 (2015). 10.1038/ncomms8455

9 Iordanescu, L., Grabowecky, M., Franconeri, S., Theeuwes, J. & Suzuki, S. Characteristic sounds make you look at target objects more quickly. Atten Percept Psychophys 72, 1736–1741 (2010). 10.3758/APP.72.7.1736

10 Janata, P. When music tells a story. Nat Neurosci 7, 203–204 (2004). 10.1038/nn0304-203

11 Ahveninen, J. et al. Evidence for distinct human auditory cortex regions for sound location versus identity processing. Nat Commun 4, 2585 (2013). 10.1038/ncomms3585

12 Rao, S. M., Mayer, A. R. & Harrington, D. L. The evolution of brain activation during temporal processing. Nat Neurosci 4, 317–323 (2001). 10.1038/85191

13 Sheth, S. A. et al. Human dorsal anterior cingulate cortex neurons mediate ongoing behavioural adaptation. Nature 488, 218–221 (2012). 10.1038/nature11239

14 Kim, C. H. et al. Melody effects on ERANm elicited by harmonic irregularity in musical syntax. Brain Res 1560, 36–45 (2014). 10.1016/j.brainres.2014.02.045

15 Luck, S. J. et al. The speed of visual attention in schizophrenia: electrophysiological and behavioral evidence. Schizophr Res 85, 174–195 (2006). 10.1016/j.schres.2006.03.040

16 Kaiser, S. et al. Intra-individual reaction time variability in schizophrenia, depression and borderline personality disorder. Brain Cogn 66, 73–82 (2008). 10.1016/j.bandc.2007.05.007

17 Binder, J. R., Liebenthal, E., Possing, E. T., Medler, D. A. & Ward, B. D. Neural correlates of sensory and decision processes in auditory object identification. Nat Neurosci 7, 295–301 (2004). 10.1038/nn1198

18 Lo, C. C. & Wang, X. J. Cortico-basal ganglia circuit mechanism for a decision threshold in reaction time tasks. Nat Neurosci 9, 956–963 (2006). 10.1038/nn1722

19 Wig, G. S., Grafton, S. T., Demos, K. E. & Kelley, W. M. Reductions in neural activity underlie behavioral components of repetition priming. Nat Neurosci 8, 1228–1233 (2005). 10.1038/nn1515

20 Stefanics, G. et al. Phase Entrainment of Human Delta Oscillations Can Mediate the Effects of Expectation on Reaction Speed. Journal of Neuroscience 30, 13578–13585 (2010). 10.1523/Jneurosci.0703-10.2010

21 Khamechian, M. B., Kozyrev, V., Treue, S., Esghaei, M. & Daliri, M. R. Routing information flow by separate neural synchrony frequencies allows for “functionally labeled lines” in higher primate cortex. Proc Natl Acad Sci U S A 116, 12506–12515 (2019). 10.1073/pnas.1819827116

22 Etzel, B. C. & Wright, E. S. Effects of Delayed Reinforcement on Response Latency and Acquisition Learning under Simultaneous and Successive Discrimination-Learning in Children. Journal of Experimental Child Psychology 1, 281–293 (1964). 10.1016/0022-0965(64)90043-8

23 Mehraei, G. et al. Auditory Brainstem Response Latency in Noise as a Marker of Cochlear Synaptopathy. Journal of Neuroscience 36, 3755–3764 (2016). 10.1523/Jneurosci.4460-15.2016

24 Bale, M. R. et al. Learning and recognition of tactile temporal sequences by mice and humans. Elife 6 (2017). 10.7554/eLife.27333

25 Esteves, M., Moreira, P. S., Sousa, N. & Leite-Almeida, H. Assessing impulsivity in humans and rodents: taking the translational road. Frontiers in Behavioral Neuroscience 15, 647922 (2021).

26 Johnson, S. A. et al. Rodent age-related impairments in discriminating perceptually similar objects parallel those observed in humans. Hippocampus 27, 759–776 (2017).

27 Koelsch, S., Schmidt, B. H. & Kansok, J. Effects of musical expertise on the early right anterior negativity: an event-related brain potential study. Psychophysiology 39, 657–663 (2002). 10.1017.S0048577202010508

28 Poulton, E. C. Perceptual Anticipation and Reaction Time. Quarterly journal of experimental psychology 2, 99–112 (1950). 10.1080/17470215008416582

29 Butz, M. V. & Hoffmann, J. Anticipations control behavior: Animal behavior in an anticipatory learning classifier system. (2002).

30 Altieri, N. & Townsend, J. T. An assessment of behavioral dynamic information processing measures in audiovisual speech perception. Front Psychol 2, 238 (2011). 10.3389/fpsyg.2011.00238

31 Eshel, N. & Roiser, J. P. Reward and punishment processing in depression. Biological psychiatry 68, 118–124 (2010). 10.1016/j.biopsych.2010.01.027

32 Kim, C. H. et al. Dissociation of Connectivity for Syntactic Irregularity and Perceptual Ambiguity in Musical Chord Stimuli. Front Neurosci 15, 693629 (2021). 10.3389/fnins.2021.693629

33 Brewer, G. A. Analyzing response time distributions. Zeitschrift für Psychologie (2015).

34 Grice, G. R., Nullmeyer, R. & Spiker, V. A. Human Reaction-Time - toward a General-Theory. Journal of Experimental Psychology-General 111, 135–153 (1982). 10.1037/0096-3445.111.1.135

35 Harris, J. A. & Carpenter, J. S. Response rate and reinforcement rate in Pavlovian conditioning. J Exp Psychol Anim Behav Process 37, 375–384 (2011). 10.1037/a0024554

36 Austen, J. M. & Sanderson, D. J. Cue duration determines response rate but not rate of acquisition of Pavlovian conditioning in mice. Quarterly journal of experimental psychology 73, 2026–2035 (2020). 10.1177/1747021820937696

37 Jennings, D. & Kirkpatrick, K. Interval duration effects on blocking in appetitive conditioning. Behav Processes 71, 318–329 (2006). 10.1016/j.beproc.2005.11.007

38 Groves, P. M. & Thompson, R. F. Habituation: a dual-process theory. Psychol Rev 77, 419–450 (1970). 10.1037/h0029810

39 de Boer, E. et al. Habituation and attention in the auditory system. Auditory system: Clinical and special topics, 343-389 (1976).

40 Thompson, R. F. Habituation: a history. Neurobiol Learn Mem 92, 127–134 (2009). 10.1016/j.nlm.2008.07.011

41 Cevik, M. O. Habituation, sensitization, and Pavlovian conditioning. Front Integr Neurosci 8, 13 (2014). 10.3389/fnint.2014.00013

42 McSweeney, F. K., Hinson, J. M. & Cannon, C. B. Sensitization-habituation may occur during operant conditioning. Psychological Bulletin 120, 256–271 (1996). 10.1037/0033-2909.120.2.256

43 McSweeney, F. K. & Murphy, E. S. Sensitization and habituation regulate reinforcer effectiveness. Neurobiol Learn Mem 92, 189–198 (2009). 10.1016/j.nlm.2008.07.002

44 Meltzer, D. Cs Duration and Reinforcement Schedule Effects on Conditioned Enhancement and Positive Conditioned Suppression. Bulletin of the Psychonomic Society 24, 290–293 (1986).

45 Thrailkill, E. A., Todd, T. P. & Bouton, M. E. Effects of conditioned stimulus (CS) duration, intertrial interval, and I/T ratio on appetitive Pavlovian conditioning. J Exp Psychol Anim Learn Cogn 46, 243–255 (2020). 10.1037/xan0000241

46 Delamater, A. R. & Holland, P. C. The influence of CS-US interval on several different indices of learning in appetitive conditioning. Journal of Experimental Psychology-Animal Behavioral Processes 34, 202–222 (2008). 10.1037/0097-7403.34.2.202

47 Odling-Smee, F. J. Background stimuli and the inter-stimulus interval during Pavlovian conditioning. Q J Exp Psychol 27, 387–392 (1975). 10.1080/14640747508400498

48 Dickinson, A. & Mackintosh, N. J. Classical conditioning in animals. Annu Rev Psychol 29, 587–612 (1978). 10.1146/annurev.ps.29.020178.003103

49 Millenson, J., Kehoe, E. J. & Gormezano, I. Classical conditioning of the rabbit’s nictitating membrane response under fixed and mixed CS-US intervals. Learning and motivation 8, 351–366 (1977).

50 Cooper, L. D. Temporal Factors in Classical-Conditioning. Learning and Motivation 22, 129–152 (1991). 10.1016/0023-9690(91)90020-9

51 Velazquez-Sanchez, C. et al. High trait impulsivity predicts food addiction-like behavior in the rat. Neuropsychopharmacology 39, 2463–2472 (2014). 10.1038/npp.2014.98

52 Celma-Miralles, A. & Toro, J. M. Non-human animals detect the rhythmic structure of a familiar tune. Psychonomic bulletin & review 27, 694–699 (2020). 10.3758/s13423-020-01739-2

53 Johnson, A. W. Examining the influence of CS duration and US density on cue-potentiated feeding through analyses of licking microstructure. Learning and Motivation 61, 85–96 (2018). 10.1016/j.lmot.2017.07.001

54 Rosas, J. M. & Alonso, G. Temporal discrimination and forgetting of CS duration in conditioned suppression. Learning and Motivation 27, 43–57 (1996). 10.1006/lmot.1996.0003

55 Schwarz-Stevens, K. S. & Cunningham, C. L. Pavlovian conditioning of heart rate and body temperature with morphine: effects of CS duration. Behav Neurosci 107, 1039–1048 (1993). 10.1037//0735-7044.107.6.1039

56 Meltzer, D. Positive conditioned suppression after shifts in CS duration and US probability. Bulletin of the Psychonomic Society 26, 565–568 (1988).

57 Meltzer, D. & Brahlek, J. A. Conditioned suppression and conditioned enhancement with the same positive UCS: an effect of CS duration. J Exp Anal Behav 13, 67–73 (1970). 10.1901/jeab.1970.13-67

58 Holland, P. C. CS-US interval as a determinant of the form of Pavlovian appetitive conditioned responses. J Exp Psychol Anim Behav Process 6, 155–174 (1980).

59 Clark, R. E. & Squire, L. R. Classical conditioning and brain systems: the role of awareness. Science 280, 77–81 (1998). 10.1126/science.280.5360.77

60 Balsam, P. Relative time in trace conditioning. Ann N Y Acad Sci 423, 211–227 (1984). 10.1111/j.1749-6632.1984.tb23432.x

61 Prinzmetal, W., McCool, C. & Park, S. Attention: reaction time and accuracy reveal different mechanisms. J Exp Psychol Gen 134, 73–92 (2005). 10.1037/0096-3445.134.1.73

62 Coddington, L. T. & Dudman, J. T. The timing of action determines reward prediction signals in identified midbrain dopamine neurons. Nature neuroscience 21, 1563–1573 (2018).

63 Eshel, N., Tian, J., Bukwich, M. & Uchida, N. Dopamine neurons share common response function for reward prediction error. Nature neuroscience 19, 479–486 (2016).

64 Anderson, C., Von Keyserlingk, M., Lidfors, L. & Weary, D. Anticipatory behaviour in animals: A critical review. Animal Welfare 29, 231–238 (2020).

65 Tosun, T., Gur, E. & Balci, F. Mice plan decision strategies based on previously learned time intervals, locations, and probabilities. Proc Natl Acad Sci U S A 113, 787–792 (2016). 10.1073/pnas.1518316113

66 Trott, J. M., Hoffman, A. N., Zhuravka, I. & Fanselow, M. S. Conditional and unconditional components of aversively motivated freezing, flight and darting in mice. Elife 11, e75663 (2022). 10.7554/eLife.75663

67 Strickland, J. A., Austen, J. M., Sprengel, R. & Sanderson, D. J. Knockout of NMDARs in CA1 and dentate gyrus fails to impair temporal control of conditioned behavior in mice. Hippocampus 34, 126–140 (2024). 10.1002/hipo.23593

68 Kail, R. Controlled and automatic processing during mental rotation. J Exp Child Psychol 51, 337–347 (1991). 10.1016/0022-0965(91)90081-3

69 Novikov, N. A. et al. Slow and Fast Responses: Two Mechanisms of Trial Outcome Processing Revealed by EEG Oscillations. Front Hum Neurosci 11, 218 (2017). 10.3389/fnhum.2017.00218

