## Supplementary figures for "Harmonization of the fastest and densest responses embodies the humanlike reaction time of mice"

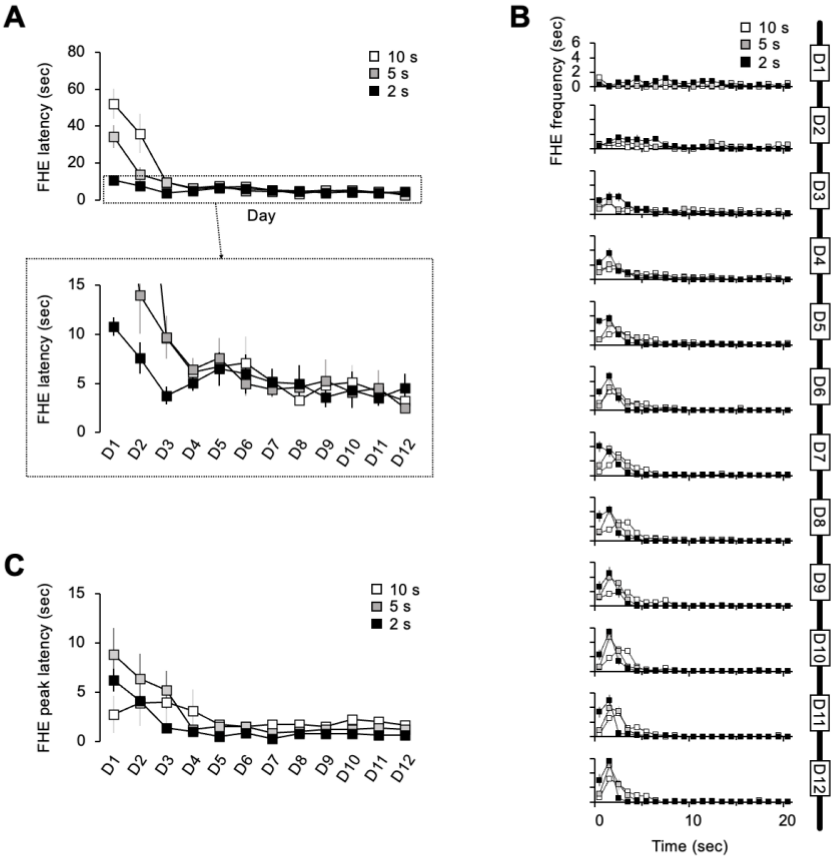


**Fig. S1. FHE and its peak. (A)** The graph depicts the FHE trace for 12 training days, calculated using the means of individual subjects' first HEs across all trials. Thus, each data point contained missing data points in which mice could not respond within the CS duration, as evidenced by the longer length of the error bar. **(B)** FHEs were extracted from the first HE in each trial. The FHE peak was extracted from a matrix containing frequency levels representing the numbers of HEs occurring within a 1-s epoch. **(C)** The FHE latency was extracted from the peak and presented in B. Error bars denote 95% confidence intervals.

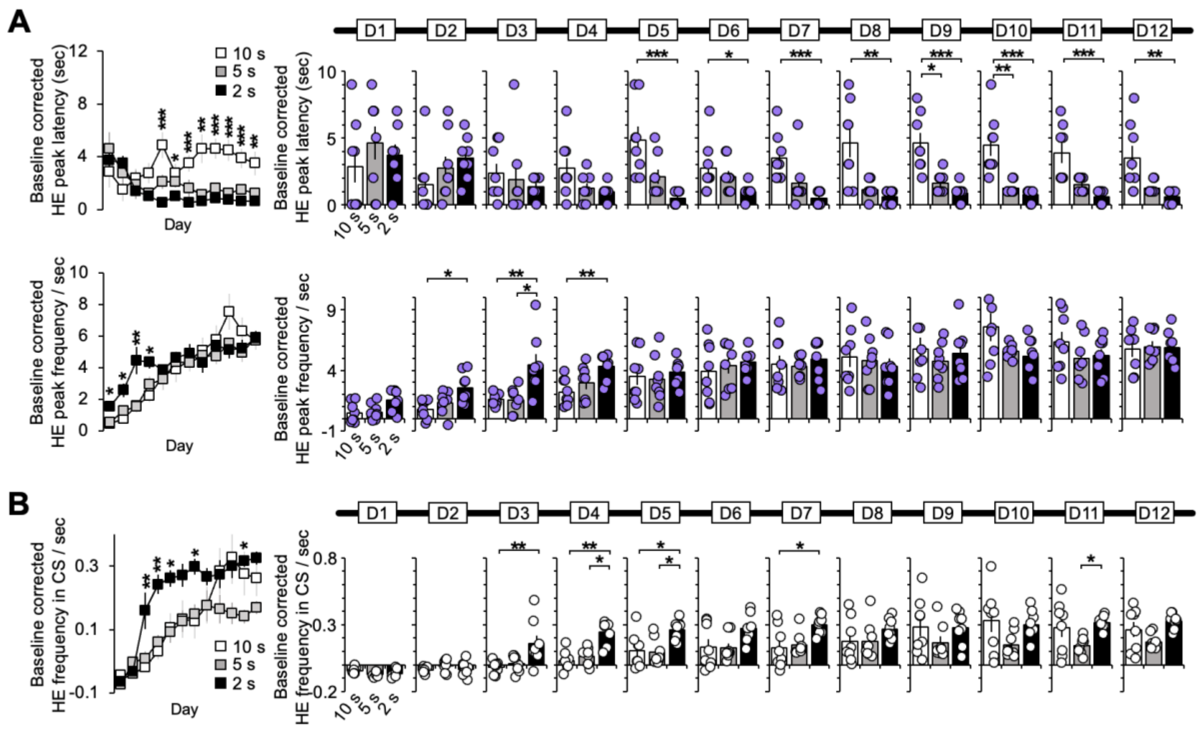


**Fig. S2. Baseline-corrected HE peak and frequency in CS.** Compared with the raw data, the peak values of HE changed for frequency but not for latency because the baseline correction procedure only affected frequency. Nevertheless, the baseline-corrected results (**A** and **B**) remained largely unchanged from the raw data. Error bars denote 95% confidence intervals. See also Tables S12 and S14 for the details of statistical analyses. *, $P < 0.05$; **, $P < 0.01$; and ***, $P < 0.001$.

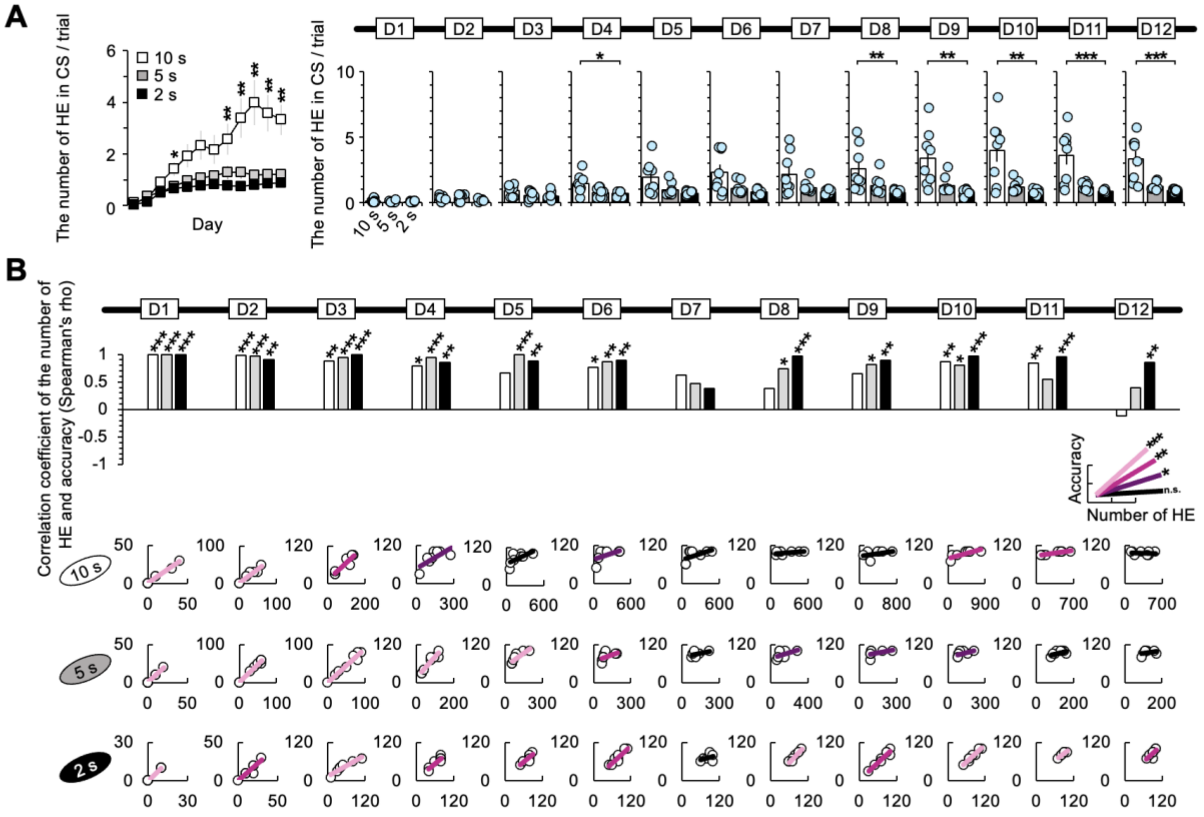


**Fig. S3. Number of HE and correlation with accuracy. (A)** The one-way ANOVA results for the number of HE revealed significant group effects on D4 and D8–D12 (Kruskal–Wallis H test, $P < 0.05$). Post hoc analysis revealed significant group differences between the 10 and 2s groups from D4 and D8–D12 (Dunn's Test, $P < 0.05$, Bonferroni-corrected). *, $P < 0.05$; **, $P < 0.01$; and ***, $P < 0.001$. Error bars denote 95% confidence intervals. **(B)** Significant positive correlations were observed across all groups (Spearman test, $P < 0.05$). For the 2s group, the significances were consistent across all the training days except for D7. The correlation test was conducted with the number of HE per trial in A, normalized to a score of 100 per HE. The nonparametric correlation was assessed using the Spearman test. See also Tables S15 and S16 for the details of statistical analyses. *, $P < 0.05$; **, $P < 0.01$; and ***, $P < 0.001$.

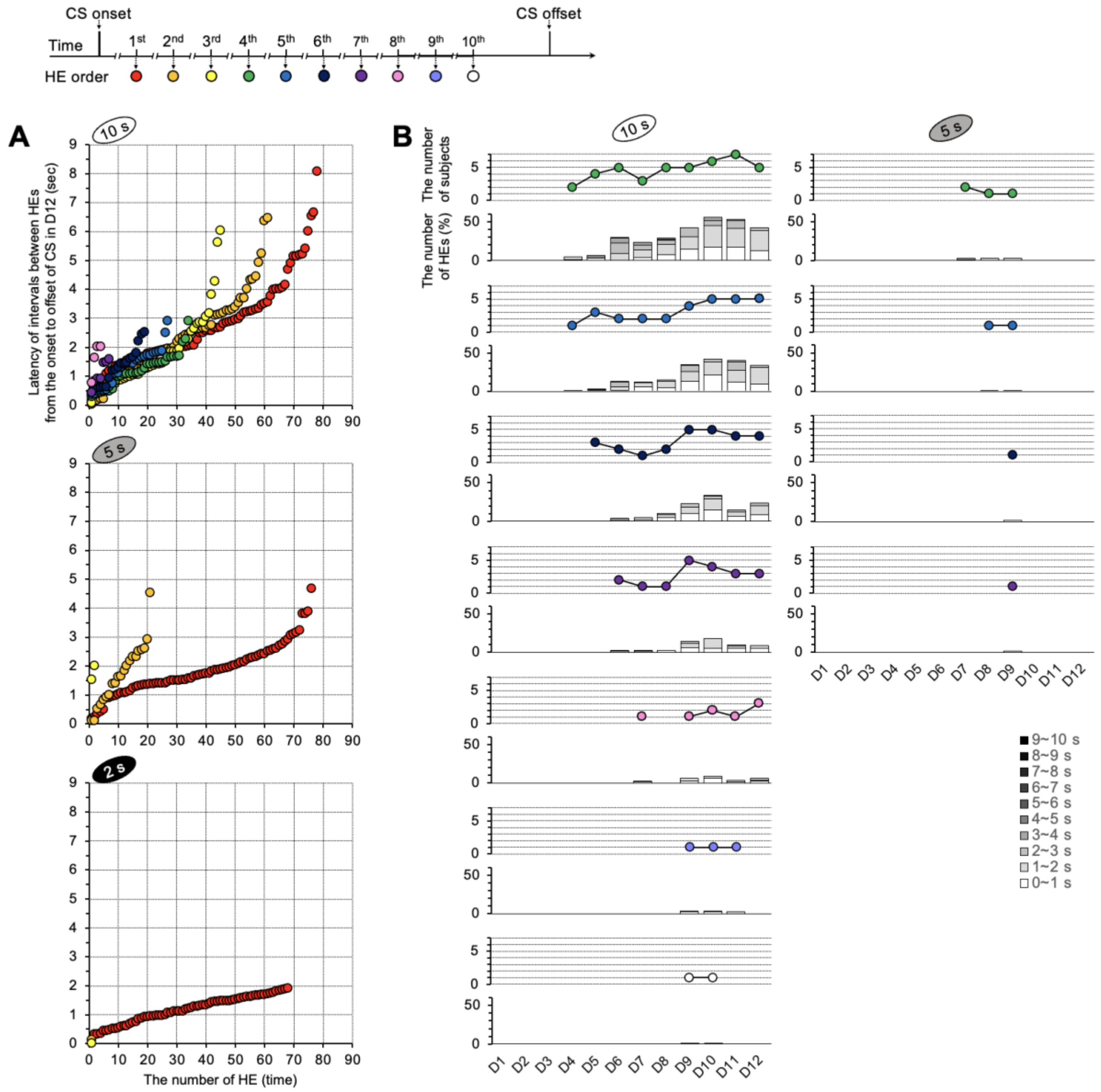


**Fig. S4. Repetition of HEs per trial. (A)** The graph depicts the latency of intervals between HEs during the CS period of D12. All HEs after the CS onset were rearranged, with the number of responses plotted on the x-axis and latency on the y-axis. The repetition of HEs in D12 was greater in the 10s group, with the maximum occurring at the eighth interval. **(B)** The number of HEs between the fourth and tenth intervals gradually dropped. Both the number of HEs and subjects were higher in the 10s than in the 5s group. Responses from the 2s group were not observed from the fourth HE onward. The majority of the eight subjects participated until the 6th HE, after which the number gradually decreased. The ninth and tenth HEs of the 10s group as well as the fifth and seventh HEs of the 5s group were observed in only one subject.

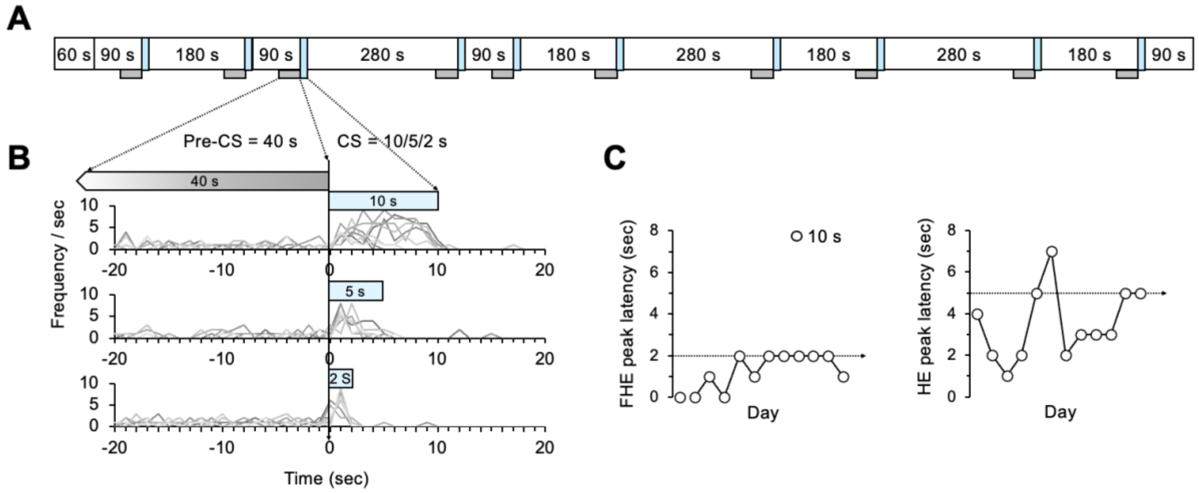


**Fig. S5. Study paradigm. (A)** Three values of intertrial interval, 90 s, 180 s, and 280 s, were randomly ordered. The CS, represented by a sky blue box, was presented between the intertrial intervals. A resting period of 60 s was included at the start of each daily experiment. The pre-CS period, based on the time window of Fig. S3B, is indicated by the gray boxes under the daily schedule. **(B)** Individual subjects' data on HE frequency were depicted as gray lines. Centered around 0, which represents the stimulus onset point, the negative section displays the pre-CS period, whereas the positive section displays the CS. **C.** The CS durations of 5 and 2s, which were the FHE and HE peak latencies in preliminary data using a 10s CS, are indicated with dotted arrow lines in each plot.

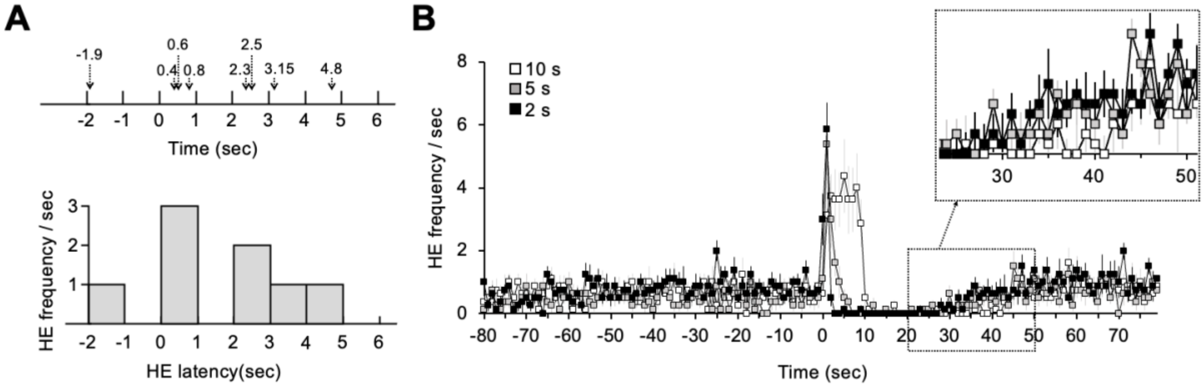


**Fig. S6. Extraction of the HE peak and pre-CS period. (A)** The occurrences of HEs within a 1-s epoch were calculated as frequency levels. The frequencies were plotted as shown in B. **(B)** HE frequency pattern in D12 shows the difference between groups after the CS onset. The time window of the pre-CS period, 40 s preceding the onset of the CS, was selected to begin after 50 s to minimize differences between the groups within the time window from 20 to 50 s after the onset of the CS, considering that the shortest intertrial interval was 90 s. The group with 10s duration for the CS had the slowest increase in frequency, most likely due to the longer duration. Error bars denote 95% confidence intervals.

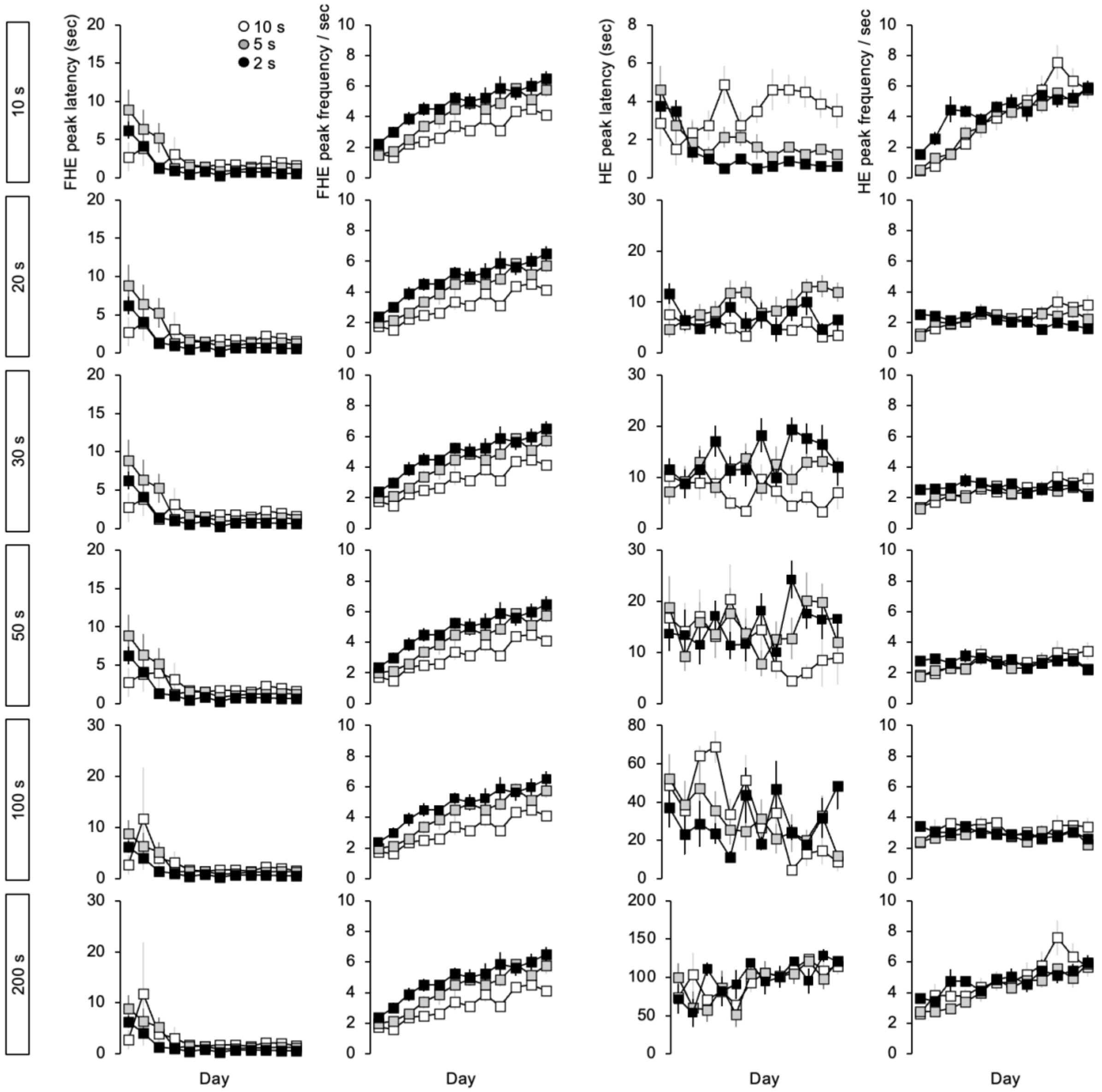


**Fig. S7. Selection of time window in the FHE and HE peaks.** We computed the FHE and HE peaks using the time windows of 10, 20, 30, 50, 100, and 200 s and then reviewed the traces. Between 20 to 100 s, there was no observed increase in the frequency of HE. In the case of 200 s, the frequency pattern demonstrated the effect of training. However, we did not select it because it was likely to include responses after the CS owing to its long duration exceeding 90 s. Finally, we selected the 10-s time window to equally estimate the FHE and HE peaks. Error bars denote 95% confidence intervals.

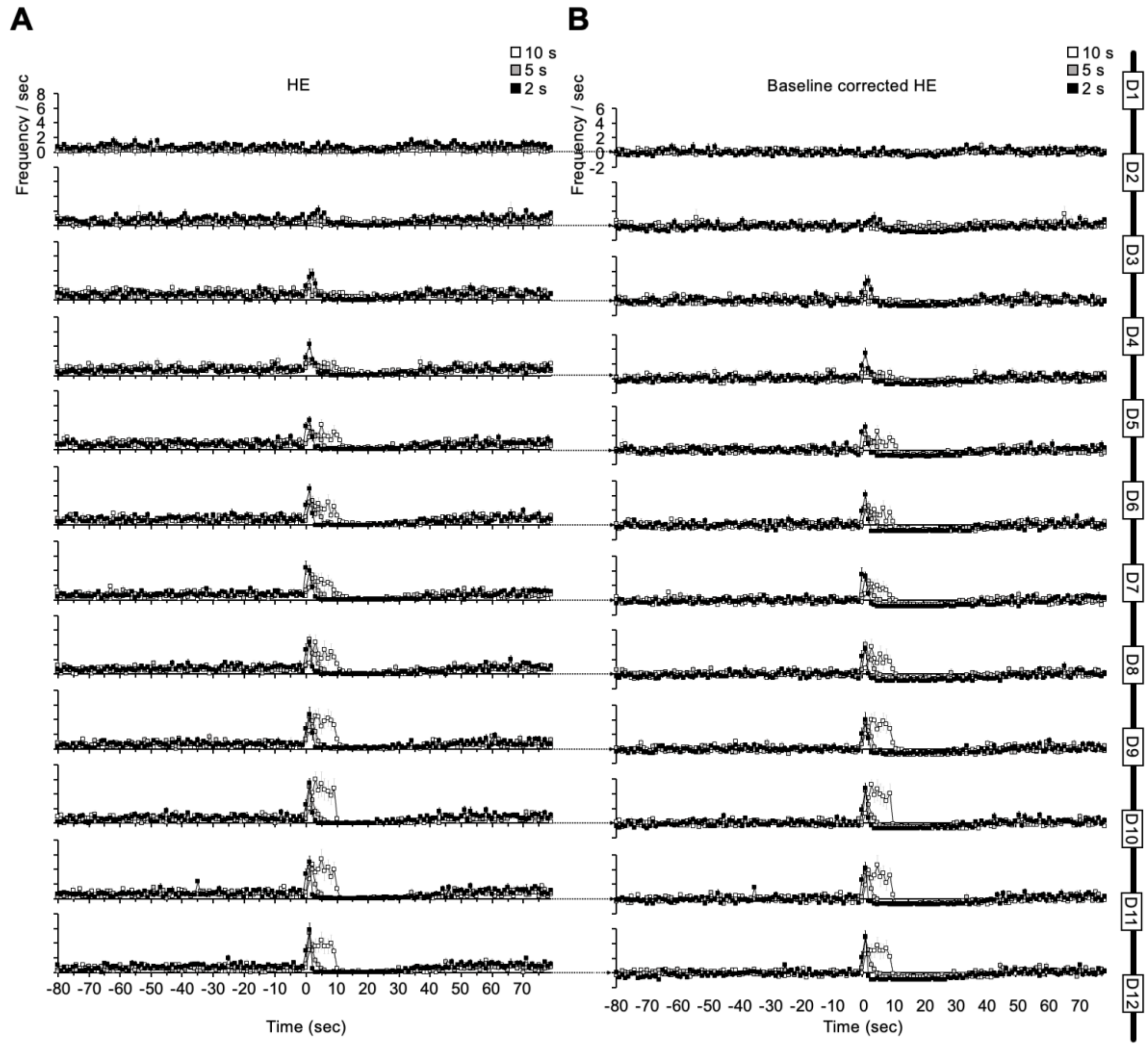


**Fig. S8. HE peak and baseline-corrected HE peak.** By subtracting the mean pre-CS value from the raw HE (**A**), we estimated the baseline-corrected HE (**B**). Error bars denote 95% confidence intervals.
