## Supplementary tables for "Harmonization of the fastest and densest responses embodies the humanlike reaction time of mice"

| Table number | Related figure | Session | Group | Component | Mean | SEM | Statistical test | Z | P | * |
|---|---|---|---|---|---|---|---|---|---|---|
| Table S1 | Fig. 2A | D1 | 10 s | FHE | 1.375 | 0.822 | Wilcoxon signed-ranks test | -1.604 | 0.109 | |
| | | | | HE | 2.875 | 1.217 | | | | |
| | | | 5 s | FHE | 4.000 | 1.225 | Wilcoxon signed-ranks test | -1 | 0.317 | |
| | | | | HE | 4.625 | 1.238 | | | | |
| | | | 2 s | FHE | 5.000 | 0.906 | Wilcoxon signed-ranks test | -1 | 0.317 | |
| | | | | HE | 3.750 | 0.773 | | | | |
| | | D2 | 10 s | FHE | 1.500 | 0.845 | Wilcoxon signed-ranks test | 0 | 1 | |
| | | | | HE | 1.500 | 0.845 | | | | |
| | | | 5 s | FHE | 2.500 | 1.118 | Wilcoxon signed-ranks test | 0 | 1 | |
| | | | | HE | 2.750 | 0.881 | | | | |
| | | | 2 s | FHE | 4.125 | 0.515 | Wilcoxon signed-ranks test | -1 | 0.317 | |
| | | | | HE | 3.500 | 0.567 | | | | |
| | | D3 | 10 s | FHE | 1.250 | 0.590 | Wilcoxon signed-ranks test | -1.342 | 0.18 | |
| | | | | HE | 2.375 | 0.706 | | | | |
| | | | 5 s | FHE | 3.375 | 1.401 | Wilcoxon signed-ranks test | -1.342 | 0.18 | |
| | | | | HE | 1.875 | 1.076 | | | | |
| | | | 2 s | FHE | 1.375 | 0.375 | Wilcoxon signed-ranks test | 0 | 1 | |
| | | | | HE | 1.375 | 0.263 | | | | |
| | | D4 | 10 s | FHE | 1.125 | 0.515 | Wilcoxon signed-ranks test | -1.476 | 0.14 | |
| | | | | HE | 2.750 | 0.773 | | | | |
| | | | 5 s | FHE | 1.250 | 0.366 | Wilcoxon signed-ranks test | 0 | 1 | |
| | | | | HE | 1.250 | 0.366 | | | | |
| | | | 2 s | FHE | 1.000 | 0.189 | Wilcoxon signed-ranks test | 0 | 1 | |
| | | | | HE | 1.000 | 0.189 | | | | |
| | | D5 | 10 s | FHE | 1.750 | 0.250 | Wilcoxon signed-ranks test | -2.207 | 0.027 | * |
| | | | | HE | 4.875 | 0.990 | | | | |
| | | | 5 s | FHE | 1.500 | 0.420 | Wilcoxon signed-ranks test | -1 | 0.317 | |
| | | | | HE | 2.130 | 0.550 | | | | |
| | | | 2 s | FHE | 0.500 | 0.189 | Wilcoxon signed-ranks test | 0 | 1 | |
| | | | | HE | 0.500 | 0.189 | | | | |
| | | D6 | 10 s | FHE | 1.500 | 0.327 | Wilcoxon signed-ranks test | -1.342 | 0.18 | |
| | | | | HE | 2.750 | 0.701 | | | | |
| | | | 5 s | FHE | 1.500 | 0.189 | Wilcoxon signed-ranks test | -1.342 | 0.18 | |
| | | | | HE | 2.125 | 0.441 | | | | |
| | | | 2 s | FHE | 0.875 | 0.227 | Wilcoxon signed-ranks test | -1 | 0.317 | |
| | | | | HE | 1.000 | 0.189 | | | | |
| | | D7 | 10 s | FHE | 1.750 | 0.164 | Wilcoxon signed-ranks test | -2.06 | 0.039 | * |
| | | | | HE | 3.500 | 0.627 | | | | |
| | | | 5 s | FHE | 0.875 | 0.227 | Wilcoxon signed-ranks test | -1 | 0.317 | |
| | | | | HE | 1.625 | 0.653 | | | | |
| | | | 2 s | FHE | 0.250 | 0.164 | Wilcoxon signed-ranks test | -1.414 | 0.157 | |
| | | | | HE | 0.500 | 0.189 | | | | |
| | | D8 | 10 s | FHE | 1.750 | 0.366 | Wilcoxon signed-ranks test | -2.041 | 0.041 | * |
| | | | | HE | 4.625 | 1.085 | | | | |
| | | | 5 s | FHE | 1.000 | 0.189 | Wilcoxon signed-ranks test | -1 | 0.317 | |
| | | | | HE | 1.125 | 0.227 | | | | |
| | | | 2 s | FHE | 0.750 | 0.250 | Wilcoxon signed-ranks test | -1 | 0.317 | |
| | | | | HE | 0.625 | 0.183 | | | | |
| | | D9 | 10 s | FHE | 1.500 | 0.535 | Wilcoxon signed-ranks test | -2.032 | 0.042 | * |
| | | | | HE | 4.625 | 0.754 | | | | |
| | | | 5 s | FHE | 1.250 | 0.250 | Wilcoxon signed-ranks test | -1 | 0.317 | |
| | | | | HE | 1.625 | 0.263 | | | | |
| | | | 2 s | FHE | 0.750 | 0.250 | Wilcoxon signed-ranks test | -1 | 0.317 | |
| | | | | HE | 0.875 | 0.227 | | | | |
| | | D10 | 10 s | FHE | 2.250 | 0.250 | Wilcoxon signed-ranks test | -1.841 | 0.066 | |
| | | | | HE | 4.500 | 0.802 | | | | |
| | | | 5 s | FHE | 1.250 | 0.164 | Wilcoxon signed-ranks test | 0 | 1 | |
| | | | | HE | 1.250 | 0.164 | | | | |
| | | | 2 s | FHE | 0.750 | 0.164 | Wilcoxon signed-ranks test | 0 | 1 | |
| | | | | HE | 0.750 | 0.164 | | | | |
| | | D11 | 10 s | FHE | 2.000 | 0.463 | Wilcoxon signed-ranks test | -2.023 | 0.043 | * |
| | | | | HE | 3.875 | 0.743 | | | | |
| | | | 5 s | FHE | 1.375 | 0.263 | Wilcoxon signed-ranks test | -1 | 0.317 | |
| | | | | HE | 1.500 | 0.189 | | | | |
| | | | 2 s | FHE | 0.625 | 0.183 | Wilcoxon signed-ranks test | 0 | 1 | |
| | | | | HE | 0.625 | 0.183 | | | | |
| | | D12 | 10 s | FHE | 1.625 | 0.263 | Wilcoxon signed-ranks test | -1.841 | 0.066 | |
| | | | | HE | 3.500 | 0.906 | | | | |
| | | | 5 s | FHE | 1.250 | 0.164 | Wilcoxon signed-ranks test | 0 | 1 | |
| | | | | HE | 1.250 | 0.164 | | | | |
| | | | 2 s | FHE | 0.630 | 0.180 | Wilcoxon signed-ranks test | 0 | 1 | |
| | | | | HE | 0.630 | 0.180 | | | | |

*P < 0.05, **P < 0.01, and ***P < 0.001

| Table number | Related figure | Session | Group | Component | Mean | SEM | Statistical test | Z | P | * |
|---|---|---|---|---|---|---|---|---|---|---|
| Table S2 | Fig. 2B | D1 | 10 s | FHE | 1.625 | 0.183 | Wilcoxon signed-ranks test | -1.734 | 0.083 | |
| | | | | HE | 0.750 | 0.313 | | | | |
| | | | 5 s | FHE | 1.500 | 0.267 | Wilcoxon signed-ranks test | -1.069 | 0.285 | |
| | | | | HE | 1.000 | 0.267 | | | | |
| | | | 2 s | FHE | 2.250 | 0.250 | Wilcoxon signed-ranks test | 0 | 1 | |
| | | | | HE | 2.250 | 0.313 | | | | |
| | | D2 | 10 s | FHE | 1.375 | 0.263 | Wilcoxon signed-ranks test | -0.577 | 0.564 | |
| | | | | HE | 1.250 | 0.313 | | | | |
| | | | 5 s | FHE | 1.750 | 0.164 | Wilcoxon signed-ranks test | -0.577 | 0.564 | |
| | | | | HE | 1.875 | 0.350 | | | | |
| | | | 2 s | FHE | 3.000 | 0.300 | Wilcoxon signed-ranks test | -1.633 | 0.102 | |
| | | | | HE | 3.500 | 0.400 | | | | |
| | | D3 | 10 s | FHE | 2.250 | 0.250 | Wilcoxon signed-ranks test | -1.414 | 0.157 | |
| | | | | HE | 2.500 | 0.189 | | | | |
| | | | 5 s | FHE | 2.500 | 0.327 | Wilcoxon signed-ranks test | -1 | 0.317 | |
| | | | | HE | 2.375 | 0.375 | | | | |
| | | | 2 s | FHE | 3.875 | 0.441 | Wilcoxon signed-ranks test | -2.07 | 0.038 | * |
| | | | | HE | 5.250 | 0.840 | | | | |
| | | D4 | 10 s | FHE | 2.375 | 0.263 | Wilcoxon signed-ranks test | -1.857 | 0.063 | |
| | | | | HE | 3.125 | 0.398 | | | | |
| | | | 5 s | FHE | 3.375 | 0.420 | Wilcoxon signed-ranks test | -1.342 | 0.18 | |
| | | | | HE | 3.750 | 0.559 | | | | |
| | | | 2 s | FHE | 4.500 | 0.423 | Wilcoxon signed-ranks test | -1.89 | 0.059 | |
| | | | | HE | 5.125 | 0.398 | | | | |
| | | D5 | 10 s | FHE | 2.625 | 0.324 | Wilcoxon signed-ranks test | -2.214 | 0.027 | * |
| | | | | HE | 4.250 | 0.726 | | | | |
| | | | 5 s | FHE | 3.875 | 0.693 | Wilcoxon signed-ranks test | -1.414 | 0.157 | |
| | | | | HE | 4.125 | 0.693 | | | | |
| | | | 2 s | FHE | 4.500 | 0.267 | Wilcoxon signed-ranks test | -1 | 0.317 | |
| | | | | HE | 4.625 | 0.324 | | | | |
| | | D6 | 10 s | FHE | 3.375 | 0.532 | Wilcoxon signed-ranks test | -1.841 | 0.066 | |
| | | | | HE | 4.750 | 0.840 | | | | |
| | | | 5 s | FHE | 4.500 | 0.707 | Wilcoxon signed-ranks test | -1.342 | 0.18 | |
| | | | | HE | 5.125 | 0.666 | | | | |
| | | | 2 s | FHE | 5.250 | 0.366 | Wilcoxon signed-ranks test | -1.414 | 0.157 | |
| | | | | HE | 5.500 | 0.327 | | | | |
| | | D7 | 10 s | FHE | 3.125 | 0.295 | Wilcoxon signed-ranks test | -2.023 | 0.043 | * |
| | | | | HE | 5.250 | 0.773 | | | | |
| | | | 5 s | FHE | 4.875 | 0.398 | Wilcoxon signed-ranks test | -1 | 0.317 | |
| | | | | HE | 5.000 | 0.327 | | | | |
| | | | 2 s | FHE | 5.000 | 0.535 | Wilcoxon signed-ranks test | -1.857 | 0.063 | |
| | | | | HE | 5.750 | 0.559 | | | | |
| | | D8 | 10 s | FHE | 3.875 | 0.479 | Wilcoxon signed-ranks test | -2.032 | 0.042 | * |
| | | | | HE | 5.750 | 0.861 | | | | |
| | | | 5 s | FHE | 4.500 | 0.463 | Wilcoxon signed-ranks test | -1.633 | 0.102 | |
| | | | | HE | 5.500 | 0.707 | | | | |
| | | | 2 s | FHE | 5.250 | 0.648 | Wilcoxon signed-ranks test | 0 | 1 | |
| | | | | HE | 5.250 | 0.648 | | | | |
| | | D9 | 10 s | FHE | 3.125 | 0.295 | Wilcoxon signed-ranks test | -2.384 | 0.017 | * |
| | | | | HE | 6.250 | 0.921 | | | | |
| | | | 5 s | FHE | 4.875 | 0.441 | Wilcoxon signed-ranks test | -1.342 | 0.180 | |
| | | | | HE | 5.500 | 0.567 | | | | |
| | | | 2 s | FHE | 5.875 | 0.766 | Wilcoxon signed-ranks test | -1.414 | 0.157 | |
| | | | | HE | 6.125 | 0.718 | | | | |
| | | D10 | 10 s | FHE | 4.375 | 0.263 | Wilcoxon signed-ranks test | -2.207 | 0.027 | * |
| | | | | HE | 8.250 | 1.176 | | | | |
| | | | 5 s | FHE | 5.875 | 0.350 | Wilcoxon signed-ranks test | -1 | 0.317 | |
| | | | | HE | 6.125 | 0.227 | | | | |
| | | | 2 s | FHE | 5.625 | 0.532 | Wilcoxon signed-ranks test | -1.414 | 0.157 | |
| | | | | HE | 5.875 | 0.515 | | | | |
| | | D11 | 10 s | FHE | 4.500 | 0.189 | Wilcoxon signed-ranks test | -2.226 | 0.026 | * |
| | | | | HE | 7.125 | 0.833 | | | | |
| | | | 5 s | FHE | 5.125 | 0.515 | Wilcoxon signed-ranks test | -2 | 0.046 | * |
| | | | | HE | 5.625 | 0.565 | | | | |
| | | | 2 s | FHE | 6.000 | 0.535 | Wilcoxon signed-ranks test | 0 | 1 | |
| | | | | HE | 6.000 | 0.535 | | | | |
| | | D12 | 10 s | FHE | 4.125 | 0.350 | Wilcoxon signed-ranks test | -2.207 | 0.027 | * |
| | | | | HE | 6.375 | 0.653 | | | | |
| | | | 5 s | FHE | 5.750 | 0.491 | Wilcoxon signed-ranks test | -1.89 | 0.059 | |
| | | | | HE | 6.375 | 0.498 | | | | |
| | | | 2 s | FHE | 6.500 | 0.500 | Wilcoxon signed-ranks test | -1 | 0.317 | |
| | | | | HE | 6.750 | 0.453 | | | | |

*P < 0.05, **P < 0.01, and ***P < 0.001

| Table number | Related figure | Session | Group | Component | Mean | Statistical test | Correlation coffeficient | P | * |
|---|---|---|---|---|---|---|---|---|---|
| Table S3 | Fig. 3A | D1 | 10 s | FHE | 1.375 | Spearman test | 0.863 | 0.006 | ** |
| | | | | HE | 2.875 | | | | |
| | | | 5 s | FHE | 4.000 | Spearman test | 0.900 | 0.002 | ** |
| | | | | HE | 4.625 | | | | |
| | | | 2 s | FHE | 5.000 | Spearman test | 0.300 | 0.470 | |
| | | | | HE | 3.750 | | | | |
| | | D2 | 10 s | FHE | 1.500 | Spearman test | 1 | 0 | *** |
| | | | | HE | 1.500 | | | | |
| | | | 5 s | FHE | 2.500 | Spearman test | -0.410 | 0.313 | |
| | | | | HE | 2.750 | | | | |
| | | | 2 s | FHE | 4.125 | Spearman test | 0.405 | 0.320 | |
| | | | | HE | 3.500 | | | | |
| | | D3 | 10 s | FHE | 1.250 | Spearman test | 0.181 | 0.669 | |
| | | | | HE | 2.375 | | | | |
| | | | 5 s | FHE | 3.375 | Spearman test | 0.433 | 0.283 | |
| | | | | HE | 1.875 | | | | |
| | | | 2 s | FHE | 1.375 | Spearman test | 0.304 | 0.463 | |
| | | | | HE | 1.375 | | | | |
| | | D4 | 10 s | FHE | 1.125 | Spearman test | -0.158 | 0.709 | |
| | | | | HE | 2.750 | | | | |
| | | | 5 s | FHE | 1.250 | Spearman test | 1 | 0 | *** |
| | | | | HE | 1.250 | | | | |
| | | | 2 s | FHE | 1.000 | Spearman test | 1 | 0 | *** |
| | | | | HE | 1.000 | | | | |
| | | D5 | 10 s | FHE | 1.750 | Spearman test | -0.033 | 0.938 | |
| | | | | HE | 4.875 | | | | |
| | | | 5 s | FHE | 1.500 | Spearman test | 0.233 | 0.579 | |
| | | | | HE | 2.130 | | | | |
| | | | 2 s | FHE | 0.500 | Spearman test | 1 | 0 | *** |
| | | | | HE | 0.500 | | | | |
| | | D6 | 10 s | FHE | 1.500 | Spearman test | -0.052 | 0.903 | |
| | | | | HE | 2.750 | | | | |
| | | | 5 s | FHE | 1.500 | Spearman test | 0.520 | 0.187 | |
| | | | | HE | 2.125 | | | | |
| | | | 2 s | FHE | 0.875 | Spearman test | 0.819 | 0.013 | * |
| | | | | HE | 1.000 | | | | |
| | | D7 | 10 s | FHE | 1.750 | Spearman test | 0 | 1 | |
| | | | | HE | 3.500 | | | | |
| | | | 5 s | FHE | 0.875 | Spearman test | 0.236 | 0.573 | |
| | | | | HE | 1.625 | | | | |
| | | | 2 s | FHE | 0.250 | Spearman test | 0.577 | 0.134 | |
| | | | | HE | 0.500 | | | | |
| | | D8 | 10 s | FHE | 1.750 | Spearman test | 0.635 | 0.090 | |
| | | | | HE | 4.625 | | | | |
| | | | 5 s | FHE | 1.000 | Spearman test | 0.819 | 0.013 | * |
| | | | | HE | 1.125 | | | | |
| | | | 2 s | FHE | 0.750 | Spearman test | 0.926 | 0.001 | ** |
| | | | | HE | 0.625 | | | | |
| | | D9 | 10 s | FHE | 1.500 | Spearman test | -0.348 | 0.398 | |
| | | | | HE | 4.625 | | | | |
| | | | 5 s | FHE | 1.250 | Spearman test | 0.200 | 0.635 | |
| | | | | HE | 1.625 | | | | |
| | | | 2 s | FHE | 0.750 | Spearman test | 0.843 | 0.009 | ** |
| | | | | HE | 0.875 | | | | |
| | | D10 | 10 s | FHE | 2.250 | Spearman test | -0.491 | 0.217 | |
| | | | | HE | 4.500 | | | | |
| | | | 5 s | FHE | 1.250 | Spearman test | 1 | 0 | *** |
| | | | | HE | 1.250 | | | | |
| | | | 2 s | FHE | 0.750 | Spearman test | 1 | 0 | *** |
| | | | | HE | 0.750 | | | | |
| | | D11 | 10 s | FHE | 2.000 | Spearman test | 0.098 | 0.818 | |
| | | | | HE | 3.875 | | | | |
| | | | 5 s | FHE | 1.375 | Spearman test | 0.956 | 0.0002 | *** |
| | | | | HE | 1.500 | | | | |
| | | | 2 s | FHE | 0.625 | Spearman test | 1 | 0 | *** |
| | | | | HE | 0.625 | | | | |
| | | D12 | 10 s | FHE | 1.625 | Spearman test | 0.462 | 0.249 | |
| | | | | HE | 3.500 | | | | |
| | | | 5 s | FHE | 1.250 | Spearman test | 1 | 0 | *** |
| | | | | HE | 1.250 | | | | |
| | | | 2 s | FHE | 0.630 | Spearman test | 1 | 0 | *** |
| | | | | HE | 0.630 | | | | |

*P < 0.05, **P < 0.01, and ***P < 0.001

| Table number | Related figure | Session | Group | Component | Mean | Statistical test | Correlation coffeficient | P | * |
|---|---|---|---|---|---|---|---|---|---|
| Table S4 | Fig. 3B | D1 | 10 s | FHE | 1.625 | Spearman test | -0.852 | 0.007 | ** |
| | | | | HE | 0.750 | | | | |
| | | | 5 s | FHE | 1.500 | Spearman test | -0.445 | 0.269 | |
| | | | | HE | 1.000 | | | | |
| | | | 2 s | FHE | 2.250 | Spearman test | 0.803 | 0.016 | * |
| | | | | HE | 2.250 | | | | |
| | | D2 | 10 s | FHE | 1.375 | Spearman test | 0.662 | 0.074 | |
| | | | | HE | 1.250 | | | | |
| | | | 5 s | FHE | 1.750 | Spearman test | 0.811 | 0.015 | * |
| | | | | HE | 1.875 | | | | |
| | | | 2 s | FHE | 3.000 | Spearman test | 0.775 | 0.024 | * |
| | | | | HE | 3.500 | | | | |
| | | D3 | 10 s | FHE | 2.250 | Spearman test | 0.777 | 0.023 | * |
| | | | | HE | 2.500 | | | | |
| | | | 5 s | FHE | 2.500 | Spearman test | 0.961 | 0.0001 | *** |
| | | | | HE | 2.375 | | | | |
| | | | 2 s | FHE | 3.875 | Spearman test | 0.614 | 0.105 | |
| | | | | HE | 5.250 | | | | |
| | | D4 | 10 s | FHE | 2.375 | Spearman test | 0.687 | 0.060 | |
| | | | | HE | 3.125 | | | | |
| | | | 5 s | FHE | 3.375 | Spearman test | 0.846 | 0.008 | ** |
| | | | | HE | 3.750 | | | | |
| | | | 2 s | FHE | 4.500 | Spearman test | 0.785 | 0.021 | * |
| | | | | HE | 5.125 | | | | |
| | | D5 | 10 s | FHE | 2.625 | Spearman test | 0.864 | 0.006 | ** |
| | | | | HE | 4.250 | | | | |
| | | | 5 s | FHE | 3.875 | Spearman test | 0.963 | 0.0001 | *** |
| | | | | HE | 4.125 | | | | |
| | | | 2 s | FHE | 4.500 | Spearman test | 0.929 | 0.0008 | *** |
| | | | | HE | 4.625 | | | | |
| | | D6 | 10 s | FHE | 3.375 | Spearman test | 0.584 | 0.129 | |
| | | | | HE | 4.750 | | | | |
| | | | 5 s | FHE | 4.500 | Spearman test | 0.723 | 0.043 | * |
| | | | | HE | 5.125 | | | | |
| | | | 2 s | FHE | 5.250 | Spearman test | 0.883 | 0.004 | ** |
| | | | | HE | 5.500 | | | | |
| | | D7 | 10 s | FHE | 3.125 | Spearman test | -0.032 | 0.940 | |
| | | | | HE | 5.250 | | | | |
| | | | 5 s | FHE | 4.875 | Spearman test | 0.981 | 0.00002 | *** |
| | | | | HE | 5.000 | | | | |
| | | | 2 s | FHE | 5.000 | Spearman test | 0.916 | 0.001 | ** |
| | | | | HE | 5.750 | | | | |
| | | D8 | 10 s | FHE | 3.875 | Spearman test | 0.415 | 0.306 | |
| | | | | HE | 5.750 | | | | |
| | | | 5 s | FHE | 4.500 | Spearman test | 0.417 | 0.304 | |
| | | | | HE | 5.500 | | | | |
| | | | 2 s | FHE | 5.250 | Spearman test | 1 | 0 | *** |
| | | | | HE | 5.250 | | | | |
| | | D9 | 10 s | FHE | 3.125 | Spearman test | 0.051 | 0.905 | |
| | | | | HE | 6.250 | | | | |
| | | | 5 s | FHE | 4.875 | Spearman test | 0.522 | 0.185 | |
| | | | | HE | 5.500 | | | | |
| | | | 2 s | FHE | 5.875 | Spearman test | 0.975 | 0.00004 | *** |
| | | | | HE | 6.125 | | | | |
| | | D10 | 10 s | FHE | 4.375 | Spearman test | -0.223 | 0.595 | |
| | | | | HE | 8.250 | | | | |
| | | | 5 s | FHE | 5.875 | Spearman test | 0.841 | 0.009 | ** |
| | | | | HE | 6.125 | | | | |
| | | | 2 s | FHE | 5.625 | Spearman test | 0.975 | 0 | *** |
| | | | | HE | 5.875 | | | | |
| | | D11 | 10 s | FHE | 4.500 | Spearman test | 0.444 | 0.270 | |
| | | | | HE | 7.125 | | | | |
| | | | 5 s | FHE | 5.125 | Spearman test | 0.950 | 0.0003 | *** |
| | | | | HE | 5.625 | | | | |
| | | | 2 s | FHE | 6.000 | Spearman test | 1 | 0 | *** |
| | | | | HE | 6.000 | | | | |
| | | D12 | 10 s | FHE | 4.125 | Spearman test | 0.119 | 0.780 | |
| | | | | HE | 6.375 | | | | |
| | | | 5 s | FHE | 5.750 | Spearman test | 0.909 | 0.002 | ** |
| | | | | HE | 6.375 | | | | |
| | | | 2 s | FHE | 6.500 | Spearman test | 0.753 | 0.031 | * |
| | | | | HE | 6.750 | | | | |

*$P < 0.05$, **$P < 0.01$, and ***$P < 0.001$

| Table number | Related figure | Session | Group | Mean | SEM | Statistical test | Test Statistic | df | P | * | post-hoc test | Post hoc pair | Test Statistic | Bonferroni corrected P | * |
|---|---|---|---|---|---|---|---|---|---|---|---|---|---|---|---|
| Table S5 | Fig. 4A | D1 | 10 s | 1.38 | 0.82 | Kruskal–Wallis H test | 6.362 | 2 | 0.042 | * | Dunn's Test | 10 s vs. 5 s | -6.125 | 0.235 | |
| | | | 5 s | 4.00 | 1.22 | | | | | | | 10 s vs. 2 s | -8.500 | 0.044 | * |
| | | | 2 s | 5.00 | 0.91 | | | | | | | 5 s vs. 2 s | -2.375 | 1.000 | |
| | | D2 | 10 s | 1.50 | 0.85 | Kruskal–Wallis H test | 8.119 | 2 | 0.017 | * | Dunn's Test | 10 s vs. 5 s | -2.875 | 1.000 | |
| | | | 5 s | 2.50 | 1.12 | | | | | | | 10 s vs. 2 s | -9.688 | 0.017 | * |
| | | | 2 s | 4.13 | 0.52 | | | | | | | 5 s vs. 2 s | -6.813 | 0.153 | |
| | | D3 | 10 s | 1.25 | 0.59 | Kruskal–Wallis H test | 1.485 | 2 | 0.476 | | Dunn's Test | 10 s vs. 5 s | | | |
| | | | 5 s | 3.38 | 1.40 | | | | | | | 10 s vs. 2 s | | | |
| | | | 2 s | 1.38 | 0.38 | | | | | | | 5 s vs. 2 s | | | |
| | | D4 | 10 s | 1.13 | 0.52 | Kruskal–Wallis H test | 0.362 | 2 | 0.835 | | Dunn's Test | 10 s vs. 5 s | | | |
| | | | 5 s | 1.25 | 0.37 | | | | | | | 10 s vs. 2 s | | | |
| | | | 2 s | 1.00 | 0.19 | | | | | | | 5 s vs. 2 s | | | |
| | | D5 | 10 s | 1.75 | 0.25 | Kruskal–Wallis H test | 0.834583901 | 2 | 0.011 | * | Dunn's Test | 10 s vs. 5 s | 6.750 | 0.126 | |
| | | | 5 s | 1.50 | 0.42 | | | | | | | 10 s vs. 2 s | 9.750 | 0.010 | * |
| | | | 2 s | 0.50 | 0.19 | | | | | | | 5 s vs. 2 s | 3.000 | 1.000 | |
| | | D6 | 10 s | 1.50 | 0.33 | Kruskal–Wallis H test | 3.78 | 2 | 0.151 | | Dunn's Test | 10 s vs. 5 s | | | |
| | | | 5 s | 1.50 | 0.19 | | | | | | | 10 s vs. 2 s | | | |
| | | | 2 s | 0.88 | 0.23 | | | | | | | 5 s vs. 2 s | | | |
| | | D7 | 10 s | 1.75 | 0.16 | Kruskal–Wallis H test | 13.918 | 2 | 0.0009 | *** | Dunn's Test | 10 s vs. 5 s | 5.250 | 0.345 | |
| | | | 5 s | 0.88 | 0.23 | | | | | | | 10 s vs. 2 s | 12.375 | 0.0006 | *** |
| | | | 2 s | 0.25 | 0.16 | | | | | | | 5 s vs. 2 s | 7.125 | 0.097 | |
| | | D8 | 10 s | 1.75 | 0.37 | Kruskal–Wallis H test | 5.182 | 2 | 0.075 | | Dunn's Test | 10 s vs. 5 s | | | |
| | | | 5 s | 1.00 | 0.19 | | | | | | | 10 s vs. 2 s | | | |
| | | | 2 s | 0.75 | 0.25 | | | | | | | 5 s vs. 2 s | | | |
| | | D9 | 10 s | 1.50 | 0.53 | Kruskal–Wallis H test | 1.735 | 2 | 0.420 | | Dunn's Test | 10 s vs. 5 s | | | |
| | | | 5 s | 1.25 | 0.25 | | | | | | | 10 s vs. 2 s | | | |
| | | | 2 s | 0.75 | 0.25 | | | | | | | 5 s vs. 2 s | | | |
| | | D10 | 10 s | 2.25 | 0.25 | Kruskal–Wallis H test | 14.054 | 2 | 0.0009 | *** | Dunn's Test | 10 s vs. 5 s | 4.250 | 0.557 | |
| | | | 5 s | 1.25 | 0.16 | | | | | | | 10 s vs. 2 s | 11.875 | 0.0006 | *** |
| | | | 2 s | 0.75 | 0.16 | | | | | | | 5 s vs. 2 s | 7.625 | 0.053 | |
| | | D11 | 10 s | 2.00 | 0.46 | Kruskal–Wallis H test | 8.514 | 2 | 0.014 | * | Dunn's Test | 10 s vs. 5 s | 6.625 | 0.131 | |
| | | | 5 s | 1.38 | 0.26 | | | | | | | 10 s vs. 2 s | 9.313 | 0.014 | * |
| | | | 2 s | 0.63 | 0.18 | | | | | | | 5 s vs. 2 s | 2.688 | 1.000 | |
| | | D12 | 10 s | 1.63 | 0.26 | Kruskal–Wallis H test | 8.531 | 2 | 0.014 | * | Dunn's Test | 10 s vs. 5 s | 5.875 | 0.163 | |
| | | | 5 s | 1.25 | 0.16 | | | | | | | 10 s vs. 2 s | 8.750 | 0.013 | * |
| | | | 2 s | 0.63 | 0.18 | | | | | | | 5 s vs. 2 s | 2.875 | 1.000 | |

*P < 0.05, **P < 0.01, and ***P < 0.001

| Table number | Related figure | Session | Group | Mean | SEM | Statistical test | Test Statistic | df | P | * | post-hoc test | Post hoc pair | Test Statistic | Bonferroni corrected P | * |
|---|---|---|---|---|---|---|---|---|---|---|---|---|---|---|---|
| Table S6 | Fig. 4A | D1 | 10 s | 1.63 | 0.18 | Kruskal–Wallis H test | 4.902 | 2 | 0.086 | | Dunn's Test | 10 s vs. 5 s | | | |
| | | | 5 s | 1.50 | 0.27 | | | | | | | 10 s vs. 2 s | | | |
| | | | 2 s | 2.25 | 0.25 | | | | | | | 5 s vs. 2 s | | | |
| | | D2 | 10 s | 1.38 | 0.26 | Kruskal–Wallis H test | 11.500 | 2 | 0.003 | ** | Dunn's Test | 10 s vs. 5 s | -2.625 | 1.000 | |
| | | | 5 s | 1.75 | 0.16 | | | | | | | 10 s vs. 2 s | -10.500 | 0.003 | ** |
| | | | 2 s | 3.00 | 0.33 | | | | | | | 5 s vs. 2 s | -7.875 | 0.044 | * |
| | | D3 | 10 s | 2.25 | 0.25 | Kruskal–Wallis H test | 8.138 | 2 | 0.017 | * | Dunn's Test | 10 s vs. 5 s | -1.750 | 1.000 | |
| | | | 5 s | 2.50 | 0.33 | | | | | | | 10 s vs. 2 s | -9.125 | 0.022 | * |
| | | | 2 s | 3.88 | 0.44 | | | | | | | 5 s vs. 2 s | -7.375 | 0.090 | |
| | | D4 | 10 s | 2.38 | 0.26 | Kruskal–Wallis H test | 10.054 | 2 | 0.007 | ** | Dunn's Test | 10 s vs. 5 s | -5.375 | 0.339 | |
| | | | 5 s | 3.38 | 0.42 | | | | | | | 10 s vs. 2 s | -10.750 | 0.005 | ** |
| | | | 2 s | 4.50 | 0.42 | | | | | | | 5 s vs. 2 s | -5.375 | 0.339 | |
| | | D5 | 10 s | 2.63 | 0.32 | Kruskal–Wallis H test | 8.981 | 2 | 0.011 | * | Dunn's Test | 10 s vs. 5 s | -5.063 | 0.424 | |
| | | | 5 s | 3.88 | 0.69 | | | | | | | 10 s vs. 2 s | -10.313 | 0.008 | ** |
| | | | 2 s | 4.50 | 0.27 | | | | | | | 5 s vs. 2 s | -5.250 | 0.381 | |
| | | D6 | 10 s | 3.38 | 0.53 | Kruskal–Wallis H test | 5.013 | 2 | 0.082 | | Dunn's Test | 10 s vs. 5 s | | | |
| | | | 5 s | 4.50 | 0.71 | | | | | | | 10 s vs. 2 s | | | |
| | | | 2 s | 5.25 | 0.37 | | | | | | | 5 s vs. 2 s | | | |
| | | D7 | 10 s | 3.13 | 0.30 | Kruskal–Wallis H test | 8.927 | 2 | 0.012 | * | Dunn's Test | 10 s vs. 5 s | -8.875 | 0.030 | * |
| | | | 5 s | 4.88 | 0.40 | | | | | | | 10 s vs. 2 s | -8.938 | 0.028 | * |
| | | | 2 s | 5.00 | 0.53 | | | | | | | 5 s vs. 2 s | -0.063 | 1.000 | |
| | | D8 | 10 s | 3.88 | 0.48 | Kruskal–Wallis H test | 2.241 | 2 | 0.326 | | Dunn's Test | 10 s vs. 5 s | | | |
| | | | 5 s | 4.50 | 0.46 | | | | | | | 10 s vs. 2 s | | | |
| | | | 2 s | 5.25 | 0.65 | | | | | | | 5 s vs. 2 s | | | |
| | | D9 | 10 s | 3.13 | 0.30 | Kruskal–Wallis H test | 10.405 | 2 | 0.006 | ** | Dunn's Test | 10 s vs. 5 s | -8.250 | 0.053 | |
| | | | 5 s | 4.88 | 0.44 | | | | | | | 10 s vs. 2 s | -10.688 | 0.006 | ** |
| | | | 2 s | 5.88 | 0.77 | | | | | | | 5 s vs. 2 s | -2.438 | 1.000 | |
| | | D10 | 10 s | 4.38 | 0.26 | Kruskal–Wallis H test | 6.904 | 2 | 0.032 | * | Dunn's Test | 10 s vs. 5 s | -7.188 | 0.112 | |
| | | | 5 s | 5.88 | 0.35 | | | | | | | 10 s vs. 2 s | -8.375 | 0.046 | * |
| | | | 2 s | 5.63 | 0.53 | | | | | | | 5 s vs. 2 s | 1.188 | 1.000 | |
| | | D11 | 10 s | 4.50 | 0.19 | Kruskal–Wallis H test | 4.017 | 2 | 0.134 | | Dunn's Test | 10 s vs. 5 s | | | |
| | | | 5 s | 5.13 | 0.52 | | | | | | | 10 s vs. 2 s | | | |
| | | | 2 s | 6.00 | 0.53 | | | | | | | 5 s vs. 2 s | | | |
| | | D12 | 10 s | 4.13 | 0.35 | Kruskal–Wallis H test | 9.962 | 2 | 0.007 | ** | Dunn's Test | 10 s vs. 5 s | -7.625 | 0.082 | |
| | | | 5 s | 5.75 | 0.49 | | | | | | | 10 s vs. 2 s | -10.563 | 0.007 | ** |
| | | | 2 s | 6.50 | 0.50 | | | | | | | 5 s vs. 2 s | -2.938 | 1.000 | |

*P < 0.05, **P < 0.01, and ***P < 0.001

| Table number | Related figure | Session | Group | Mean | SEM | Statistical test | Test Statistic | df | P | * | post-hoc test | Post hoc pair | Test Statistic | Bonferroni corrected P | * |
|---|---|---|---|---|---|---|---|---|---|---|---|---|---|---|---|
| Table S7 | Fig. 4B | D1 | 10 s | 2.88 | 1.22 | Kruskal–Wallis H test | 1.549 | 2 | 0.461 | | Dunn's Test | 10 s vs. 5 s | | | |
| | | | 5 s | 4.63 | 1.24 | | | | | | | 10 s vs. 2 s | | | |
| | | | 2 s | 3.75 | 0.77 | | | | | | | 5 s vs. 2 s | | | |
| | | D2 | 10 s | 1.50 | 0.85 | Kruskal–Wallis H test | 5.122 | 2 | 0.077 | | Dunn's Test | 10 s vs. 5 s | | | |
| | | | 5 s | 2.75 | 0.88 | | | | | | | 10 s vs. 2 s | | | |
| | | | 2 s | 3.50 | 0.57 | | | | | | | 5 s vs. 2 s | | | |
| | | D3 | 10 s | 2.38 | 0.71 | Kruskal–Wallis H test | 1.468 | 2 | 0.480 | | Dunn's Test | 10 s vs. 5 s | | | |
| | | | 5 s | 1.88 | 1.08 | | | | | | | 10 s vs. 2 s | | | |
| | | | 2 s | 1.38 | 0.26 | | | | | | | 5 s vs. 2 s | | | |
| | | D4 | 10 s | 2.75 | 0.77 | Kruskal–Wallis H test | 4.849 | 2 | 0.089 | | Dunn's Test | 10 s vs. 5 s | | | |
| | | | 5 s | 1.25 | 0.37 | | | | | | | 10 s vs. 2 s | | | |
| | | | 2 s | 1.00 | 0.19 | | | | | | | 5 s vs. 2 s | | | |
| | | D5 | 10 s | 4.88 | 0.99 | Kruskal–Wallis H test | 15.765 | 2 | 0.0004 | *** | Dunn's Test | 10 s vs. 5 s | 7.313 | 0.102 | |
| | | | 5 s | 2.13 | 0.55 | | | | | | | 10 s vs. 2 s | 13.688 | 0.0002 | *** |
| | | | 2 s | 0.50 | 0.19 | | | | | | | 5 s vs. 2 s | 6.375 | 0.194 | |
| | | D6 | 10 s | 2.75 | 0.70 | Kruskal–Wallis H test | 7.498 | 2 | 0.024 | * | Dunn's Test | 10 s vs. 5 s | 6.750 | 0.125 | |
| | | | 5 s | 2.13 | 0.44 | | | | | | | 10 s vs. 2 s | 8.625 | 0.028 | * |
| | | | 2 s | 1.00 | 0.19 | | | | | | | 5 s vs. 2 s | 1.875 | 1.000 | |
| | | D7 | 10 s | 3.50 | 0.63 | Kruskal–Wallis H test | 14.557 | 2 | 0.0007 | *** | Dunn's Test | 10 s vs. 5 s | 5.063 | 0.416 | |
| | | | 5 s | 1.63 | 0.65 | | | | | | | 10 s vs. 2 s | 12.938 | 0.0005 | *** |
| | | | 2 s | 0.50 | 0.19 | | | | | | | 5 s vs. 2 s | 7.875 | 0.064 | |
| | | D8 | 10 s | 4.63 | 1.08 | Kruskal–Wallis H test | 12.069 | 2 | 0.002 | ** | Dunn's Test | 10 s vs. 5 s | 3.750 | 0.766 | |
| | | | 5 s | 1.13 | 0.23 | | | | | | | 10 s vs. 2 s | 11.250 | 0.002 | ** |
| | | | 2 s | 0.63 | 0.18 | | | | | | | 5 s vs. 2 s | 7.500 | 0.069 | |
| | | D9 | 10 s | 4.63 | 0.75 | Kruskal–Wallis H test | 16.145 | 2 | 0.0003 | *** | Dunn's Test | 10 s vs. 5 s | 4.500 | 0.565 | |
| | | | 5 s | 1.63 | 0.26 | | | | | | | 10 s vs. 2 s | 13.500 | 0.0002 | *** |
| | | | 2 s | 0.88 | 0.23 | | | | | | | 5 s vs. 2 s | 9.000 | 0.026 | * |
| | | D10 | 10 s | 4.50 | 0.80 | Kruskal–Wallis H test | 18.289 | 2 | 0.0001 | *** | Dunn's Test | 10 s vs. 5 s | 3.625 | 0.816 | |
| | | | 5 s | 1.25 | 0.16 | | | | | | | 10 s vs. 2 s | 13.625 | 0.0001 | *** |
| | | | 2 s | 0.75 | 0.16 | | | | | | | 5 s vs. 2 s | 10.000 | 0.007 | ** |
| | | D11 | 10 s | 3.88 | 0.74 | Kruskal–Wallis H test | 16.653 | 2 | 0.0002 | *** | Dunn's Test | 10 s vs. 5 s | 6.500 | 0.162 | |
| | | | 5 s | 1.50 | 0.19 | | | | | | | 10 s vs. 2 s | 13.750 | 0.0001 | *** |
| | | | 2 s | 0.63 | 0.18 | | | | | | | 5 s vs. 2 s | 7.250 | 0.095 | |
| | | D12 | 10 s | 3.50 | 0.91 | Kruskal–Wallis H test | 12.427 | 2 | 0.002 | ** | Dunn's Test | 10 s vs. 5 s | 5.125 | 0.338 | |
| | | | 5 s | 1.25 | 0.16 | | | | | | | 10 s vs. 2 s | 11.375 | 0.0013 | ** |
| | | | 2 s | 0.63 | 0.18 | | | | | | | 5 s vs. 2 s | 6.250 | 0.159 | |

*P < 0.05, **P < 0.01, and ***P < 0.001

| Table number | Related figure | Session | Group | Mean | SEM | Statistical test | Test Statistic | df | P | * | post-hoc test | Post hoc pair | Test Statistic | Bonferroni corrected P | * |
|---|---|---|---|---|---|---|---|---|---|---|---|---|---|---|---|
| ***Table S8*** | ***Fig. 4B*** | ***D1*** | ***10 s*** | 0.75 | 0.31 | Kruskal–Wallis H test | 8.808 | 2 | 0.012 | * | Dunn's Test | ***10 s vs. 5 s*** | -1.750 | 1.000 | |
| | | | ***5 s*** | 1.00 | 0.27 | | | | | | | ***10 s vs. 2 s*** | -9.500 | 0.016 | * |
| | | | ***2 s*** | 2.25 | 0.31 | | | | | | | ***5 s vs. 2 s*** | -7.750 | 0.069 | |
| | | ***D2*** | ***10 s*** | 1.25 | 0.31 | Kruskal–Wallis H test | 11.410 | 2 | 0.003 | ** | Dunn's Test | ***10 s vs. 5 s*** | -3.750 | 0.806 | |
| | | | ***5 s*** | 1.88 | 0.35 | | | | | | | ***10 s vs. 2 s*** | -11.250 | 0.003 | ** |
| | | | ***2 s*** | 3.50 | 0.42 | | | | | | | ***5 s vs. 2 s*** | -7.500 | 0.081 | |
| | | ***D3*** | ***10 s*** | 2.50 | 0.19 | Kruskal–Wallis H test | 10.724 | 2 | 0.005 | ** | Dunn's Test | ***10 s vs. 5 s*** | 0.500 | 1.000 | |
| | | | ***5 s*** | 2.38 | 0.38 | | | | | | | ***10 s vs. 2 s*** | -10.000 | 0.011 | ** |
| | | | ***2 s*** | 5.25 | 0.84 | | | | | | | ***5 s vs. 2 s*** | -9.500 | 0.017 | ** |
| | | ***D4*** | ***10 s*** | 3.13 | 0.40 | Kruskal–Wallis H test | 7.604 | 2 | 0.022 | * | Dunn's Test | ***10 s vs. 5 s*** | -2.875 | 1.000 | |
| | | | ***5 s*** | 3.75 | 0.56 | | | | | | | ***10 s vs. 2 s*** | -9.313 | 0.021 | * |
| | | | ***2 s*** | 5.13 | 0.40 | | | | | | | ***5 s vs. 2 s*** | -6.438 | 0.188 | |
| | | ***D5*** | ***10 s*** | 4.25 | 0.73 | Kruskal–Wallis H test | 1.044 | 2 | 0.593 | | Dunn's Test | ***10 s vs. 5 s*** | | | |
| | | | ***5 s*** | 4.13 | 0.69 | | | | | | | ***10 s vs. 2 s*** | | | |
| | | | ***2 s*** | 4.63 | 0.32 | | | | | | | ***5 s vs. 2 s*** | | | |
| | | ***D6*** | ***10 s*** | 4.75 | 0.84 | Kruskal–Wallis H test | 0.361 | 2 | 0.835 | | Dunn's Test | ***10 s vs. 5 s*** | | | |
| | | | ***5 s*** | 5.13 | 0.67 | | | | | | | ***10 s vs. 2 s*** | | | |
| | | | ***2 s*** | 5.50 | 0.33 | | | | | | | ***5 s vs. 2 s*** | | | |
| | | ***D7*** | ***10 s*** | 5.25 | 0.77 | Kruskal–Wallis H test | 0.835 | 2 | 0.542 | | Dunn's Test | ***10 s vs. 5 s*** | | | |
| | | | ***5 s*** | 5.00 | 0.33 | | | | | | | ***10 s vs. 2 s*** | | | |
| | | | ***2 s*** | 5.75 | 0.56 | | | | | | | ***5 s vs. 2 s*** | | | |
| | | ***D8*** | ***10 s*** | 5.75 | 0.86 | Kruskal–Wallis H test | 0.117 | 2 | 0.943 | | Dunn's Test | ***10 s vs. 5 s*** | | | |
| | | | ***5 s*** | 5.50 | 0.71 | | | | | | | ***10 s vs. 2 s*** | | | |
| | | | ***2 s*** | 5.25 | 0.65 | | | | | | | ***5 s vs. 2 s*** | | | |
| | | ***D9*** | ***10 s*** | 6.25 | 0.92 | Kruskal–Wallis H test | 0.282 | 2 | 0.869 | | Dunn's Test | ***10 s vs. 5 s*** | | | |
| | | | ***5 s*** | 5.50 | 0.57 | | | | | | | ***10 s vs. 2 s*** | | | |
| | | | ***2 s*** | 6.13 | 0.72 | | | | | | | ***5 s vs. 2 s*** | | | |
| | | ***D10*** | ***10 s*** | 8.25 | 1.18 | Kruskal–Wallis H test | 2.834 | 2 | 0.242 | | Dunn's Test | ***10 s vs. 5 s*** | | | |
| | | | ***5 s*** | 6.13 | 0.23 | | | | | | | ***10 s vs. 2 s*** | | | |
| | | | ***2 s*** | 5.88 | 0.52 | | | | | | | ***5 s vs. 2 s*** | | | |
| | | ***D11*** | ***10 s*** | 7.13 | 0.83 | Kruskal–Wallis H test | 1.871 | 2 | 0.392 | | Dunn's Test | ***10 s vs. 5 s*** | | | |
| | | | ***5 s*** | 5.63 | 0.56 | | | | | | | ***10 s vs. 2 s*** | | | |
| | | | ***2 s*** | 6.00 | 0.53 | | | | | | | ***5 s vs. 2 s*** | | | |
| | | ***D12*** | ***10 s*** | 6.38 | 0.65 | Kruskal–Wallis H test | 0.284 | 2 | 0.868 | | Dunn's Test | ***10 s vs. 5 s*** | | | |
| | | | ***5 s*** | 6.38 | 0.50 | | | | | | | ***10 s vs. 2 s*** | | | |
| | | | ***2 s*** | 6.75 | 0.45 | | | | | | | ***5 s vs. 2 s*** | | | |

**P < 0.05, **P < 0.01, and ***P < 0.001*

| Table number | Related figure | Session | Group | Mean | SEM | Statistical test | Test Statistic | df | P | * | post-hoc test | Post hoc pair | Test Statistic | Bonferroni corrected P | * |
|---|---|---|---|---|---|---|---|---|---|---|---|---|---|---|---|
| Table S9 | Fig. 5A | D1 | 10 s | 8.75 | 3.98 | Kruskal–Wallis H test | 1.989 | 2 | 0.370 | | Dunn's Test | 10 s vs. 5 s | | | |
| | | | 5 s | 5.00 | 2.67 | | | | | | | 10 s vs. 2 s | | | |
| | | | 2 s | 3.75 | 1.83 | | | | | | | 5 s vs. 2 s | | | |
| | | D2 | 10 s | 22.50 | 5.90 | Kruskal–Wallis H test | 5.304 | 2 | 0.071 | | Dunn's Test | 10 s vs. 5 s | | | |
| | | | 5 s | 36.25 | 8.65 | | | | | | | 10 s vs. 2 s | | | |
| | | | 2 s | 13.75 | 3.24 | | | | | | | 5 s vs. 2 s | | | |
| | | D3 | 10 s | 61.25 | 8.95 | Kruskal–Wallis H test | 2.004 | 2 | 0.367 | | Dunn's Test | 10 s vs. 5 s | | | |
| | | | 5 s | 47.50 | 10.48 | | | | | | | 10 s vs. 2 s | | | |
| | | | 2 s | 42.50 | 6.48 | | | | | | | 5 s vs. 2 s | | | |
| | | D4 | 10 s | 81.25 | 8.33 | Kruskal–Wallis H test | 5.055 | 2 | 0.080 | | Dunn's Test | 10 s vs. 5 s | | | |
| | | | 5 s | 63.75 | 9.44 | | | | | | | 10 s vs. 2 s | | | |
| | | | 2 s | 60.00 | 5.98 | | | | | | | 5 s vs. 2 s | | | |
| | | D5 | 10 s | 85.00 | 6.27 | Kruskal–Wallis H test | 3.715 | 2 | 0.156 | | Dunn's Test | 10 s vs. 5 s | | | |
| | | | 5 s | 77.50 | 6.20 | | | | | | | 10 s vs. 2 s | | | |
| | | | 2 s | 68.75 | 4.79 | | | | | | | 5 s vs. 2 s | | | |
| | | D6 | 10 s | 91.25 | 6.39 | Kruskal–Wallis H test | 4.84 | 2 | 0.089 | | Dunn's Test | 10 s vs. 5 s | | | |
| | | | 5 s | 82.50 | 4.53 | | | | | | | 10 s vs. 2 s | | | |
| | | | 2 s | 72.50 | 7.01 | | | | | | | 5 s vs. 2 s | | | |
| | | D7 | 10 s | 88.75 | 6.39 | Kruskal–Wallis H test | 7.509 | 2 | 0.023 | * | Dunn's Test | 10 s vs. 5 s | -0.6875 | 1.000 | |
| | | | 5 s | 90.00 | 2.67 | | | | | | | 10 s vs. 2 s | -8.5 | 0.041 | * |
| | | | 2 s | 72.50 | 3.66 | | | | | | | 5 s vs. 2 s | -7.8125 | 0.070 | |
| | | D8 | 10 s | 97.50 | 1.64 | Kruskal–Wallis H test | 8.333 | 2 | 0.016 | * | Dunn's Test | 10 s vs. 5 s | -4.25 | 0.610 | |
| | | | 5 s | 88.75 | 4.41 | | | | | | | 10 s vs. 2 s | -9.625 | 0.012 | * |
| | | | 2 s | 76.25 | 5.65 | | | | | | | 5 s vs. 2 s | -5.375 | 0.323 | |
| | | D9 | 10 s | 93.75 | 3.75 | Kruskal–Wallis H test | 6.918 | 2 | 0.031 | * | Dunn's Test | 10 s vs. 5 s | -1.0625 | 1.000 | |
| | | | 5 s | 92.50 | 3.66 | | | | | | | 10 s vs. 2 s | -8.125 | 0.047 | * |
| | | | 2 s | 71.25 | 8.33 | | | | | | | 5 s vs. 2 s | -7.0625 | 0.106 | |
| | | D10 | 10 s | 93.75 | 3.75 | Kruskal–Wallis H test | 5.379 | 2 | 0.068 | | Dunn's Test | 10 s vs. 5 s | | | |
| | | | 5 s | 91.25 | 2.95 | | | | | | | 10 s vs. 2 s | | | |
| | | | 2 s | 76.25 | 6.25 | | | | | | | 5 s vs. 2 s | | | |
| | | D11 | 10 s | 96.25 | 1.83 | Kruskal–Wallis H test | 9.692 | 2 | 0.008 | ** | Dunn's Test | 10 s vs. 5 s | -2.0625 | 1.000 | |
| | | | 5 s | 93.75 | 2.63 | | | | | | | 10 s vs. 2 s | -9.75 | 0.009 | ** |
| | | | 2 s | 83.75 | 2.63 | | | | | | | 5 s vs. 2 s | -7.6875 | 0.060 | |
| | | D12 | 10 s | 96.25 | 1.83 | Kruskal–Wallis H test | 4.336 | 2 | 0.114 | | Dunn's Test | 10 s vs. 5 s | | | |
| | | | 5 s | 95.00 | 2.67 | | | | | | | 10 s vs. 2 s | | | |
| | | | 2 s | 86.25 | 4.20 | | | | | | | 5 s vs. 2 s | | | |

*$P < 0.05$, **$P < 0.01$, and ***$P < 0.001$

| ***Table number*** | ***Related figure*** | ***Session*** | ***Group*** | ***Mean*** | ***SEM*** | ***Statistical test*** | ***Test Statistic*** | ***df*** | ***P*** | ***** | ***post-hoc test*** | ***Post hoc pair*** | ***Test Statistic*** | ***Bonferroni corrected P*** | ***** |
|---|---|---|---|---|---|---|---|---|---|---|---|---|---|---|---|
| ***Table S10*** | ***Fig. 5B*** | ***D1*** | ***10 s*** | 0.011 | 0.005 | Kruskal–Wallis H test | 0.286 | 2 | 0.867 | | Dunn's Test | ***10 s vs. 5 s*** | | | |
| | | | ***5 s*** | 0.010 | 0.005 | | | | | | | ***10 s vs. 2 s*** | | | |
| | | | ***2 s*** | 0.019 | 0.009 | | | | | | | ***5 s vs. 2 s*** | | | |
| | | ***D2*** | ***10 s*** | 0.028 | 0.008 | Kruskal–Wallis H test | 5.546 | 2 | 0.062 | | Dunn's Test | ***10 s vs. 5 s*** | | | |
| | | | ***5 s*** | 0.075 | 0.018 | | | | | | | ***10 s vs. 2 s*** | | | |
| | | | ***2 s*** | 0.075 | 0.016 | | | | | | | ***5 s vs. 2 s*** | | | |
| | | ***D3*** | ***10 s*** | 0.090 | 0.015 | Kruskal–Wallis H test | 9.899 | 2 | 0.007 | ** | Dunn's Test | ***10 s vs. 5 s*** | -2.375 | 1.000 | |
| | | | ***5 s*** | 0.105 | 0.023 | | | | | | | ***10 s vs. 2 s*** | -10.563 | 0.008 | ** |
| | | | ***2 s*** | 0.250 | 0.055 | | | | | | | ***5 s vs. 2 s*** | -8.188 | 0.060 | |
| | | ***D4*** | ***10 s*** | 0.143 | 0.026 | Kruskal–Wallis H test | 12.39 | 2 | 0.002 | ** | Dunn's Test | ***10 s vs. 5 s*** | -0.875 | 1.000 | |
| | | | ***5 s*** | 0.150 | 0.027 | | | | | | | ***10 s vs. 2 s*** | -11.125 | 0.005 | ** |
| | | | ***2 s*** | 0.338 | 0.032 | | | | | | | ***5 s vs. 2 s*** | -10.250 | 0.011 | * |
| | | ***D5*** | ***10 s*** | 0.193 | 0.045 | Kruskal–Wallis H test | 9.224 | 2 | 0.010 | * | Dunn's Test | ***10 s vs. 5 s*** | -1.000 | 1.000 | |
| | | | ***5 s*** | 0.193 | 0.036 | | | | | | | ***10 s vs. 2 s*** | -9.688 | 0.017 | * |
| | | | ***2 s*** | 0.363 | 0.025 | | | | | | | ***5 s vs. 2 s*** | -8.688 | 0.040 | * |
| | | ***D6*** | ***10 s*** | 0.234 | 0.056 | Kruskal–Wallis H test | 6.199 | 2 | 0.045 | * | Dunn's Test | ***10 s vs. 5 s*** | -1.500 | 1.000 | |
| | | | ***5 s*** | 0.215 | 0.035 | | | | | | | ***10 s vs. 2 s*** | -6.750 | 0.168 | |
| | | | ***2 s*** | 0.394 | 0.035 | | | | | | | ***5 s vs. 2 s*** | -8.250 | 0.058 | |
| | | ***D7*** | ***10 s*** | 0.218 | 0.056 | Kruskal–Wallis H test | 9.386 | 2 | 0.009 | ** | Dunn's Test | ***10 s vs. 5 s*** | -1.438 | 1.000 | |
| | | | ***5 s*** | 0.230 | 0.033 | | | | | | | ***10 s vs. 2 s*** | -10.000 | 0.014 | * |
| | | | ***2 s*** | 0.413 | 0.026 | | | | | | | ***5 s vs. 2 s*** | -8.563 | 0.046 | * |
| | | ***D8*** | ***10 s*** | 0.256 | 0.060 | Kruskal–Wallis H test | 4.883 | 2 | 0.087 | | Dunn's Test | ***10 s vs. 5 s*** | | | |
| | | | ***5 s*** | 0.258 | 0.052 | | | | | | | ***10 s vs. 2 s*** | | | |
| | | | ***2 s*** | 0.388 | 0.026 | | | | | | | ***5 s vs. 2 s*** | | | |
| | | ***D9*** | ***10 s*** | 0.339 | 0.076 | Kruskal–Wallis H test | 1.875 | 2 | 0.392 | | Dunn's Test | ***10 s vs. 5 s*** | | | |
| | | | ***5 s*** | 0.258 | 0.047 | | | | | | | ***10 s vs. 2 s*** | | | |
| | | | ***2 s*** | 0.375 | 0.044 | | | | | | | ***5 s vs. 2 s*** | | | |
| | | ***D10*** | ***10 s*** | 0.400 | 0.087 | Kruskal–Wallis H test | 5.554 | 2 | 0.062 | | Dunn's Test | ***10 s vs. 5 s*** | | | |
| | | | ***5 s*** | 0.235 | 0.033 | | | | | | | ***10 s vs. 2 s*** | | | |
| | | | ***2 s*** | 0.400 | 0.034 | | | | | | | ***5 s vs. 2 s*** | | | |
| | | ***D11*** | ***10 s*** | 0.359 | 0.071 | Kruskal–Wallis H test | 7.196 | 2 | 0.027 | * | Dunn's Test | ***10 s vs. 5 s*** | -6.313 | 0.220 | |
| | | | ***5 s*** | 0.240 | 0.021 | | | | | | | ***10 s vs. 2 s*** | -2.938 | 1.000 | |
| | | | ***2 s*** | 0.425 | 0.016 | | | | | | | ***5 s vs. 2 s*** | -9.250 | 0.026 | * |
| | | ***D12*** | ***10 s*** | 0.333 | 0.059 | Kruskal–Wallis H test | 9.378 | 2 | 0.009 | ** | Dunn's Test | ***10 s vs. 5 s*** | -4.000 | 0.761 | |
| | | | ***5 s*** | 0.243 | 0.020 | | | | | | | ***10 s vs. 2 s*** | -6.625 | 0.176 | |
| | | | ***2 s*** | 0.444 | 0.018 | | | | | | | ***5 s vs. 2 s*** | -10.625 | 0.007 | ** |

**P < 0.05, **P < 0.01, and ***P < 0.001*

| Table number | Related figure | Session | Group | Mean | SEM | Statistical test | Test Statistic | df | P | * | post-hoc test | Post hoc pair | Test Statistic | Bonferroni corrected P | * |
|---|---|---|---|---|---|---|---|---|---|---|---|---|---|---|---|
| ***Table S11*** | ***Fig. 5B*** | ***D1*** | ***10 s*** | 0.056 | 0.003 | Kruskal–Wallis H test | 14.236 | 2 | 0.0008 | *** | Dunn's Test | ***10 s vs. 5 s*** | -11.438 | 0.004 | ** |
| | | | ***5 s*** | 0.084 | 0.003 | | | | | | | ***10 s vs. 2 s*** | -11.625 | 0.003 | ** |
| | | | ***2 s*** | 0.083 | 0.005 | | | | | | | ***5 s vs. 2 s*** | 0.188 | 1.000 | |
| | | ***D2*** | ***10 s*** | 0.067 | 0.004 | Kruskal–Wallis H test | 14.753 | 2 | 0.004 | ** | Dunn's Test | ***10 s vs. 5 s*** | -3.875 | 0.817 | |
| | | | ***5 s*** | 0.078 | 0.005 | | | | | | | ***10 s vs. 2 s*** | -13.188 | 0.001 | ** |
| | | | ***2 s*** | 0.109 | 0.005 | | | | | | | ***5 s vs. 2 s*** | -9.313 | 0.025 | * |
| | | ***D3*** | ***10 s*** | 0.110 | 0.007 | Kruskal–Wallis H test | 4.32 | 2 | 0.115 | | Dunn's Test | ***10 s vs. 5 s*** | | | |
| | | | ***5 s*** | 0.097 | 0.003 | | | | | | | ***10 s vs. 2 s*** | | | |
| | | | ***2 s*** | 0.091 | 0.006 | | | | | | | ***5 s vs. 2 s*** | | | |
| | | ***D4*** | ***10 s*** | 0.113 | 0.006 | Kruskal–Wallis H test | 4.789 | 2 | 0.091 | | Dunn's Test | ***10 s vs. 5 s*** | | | |
| | | | ***5 s*** | 0.089 | 0.007 | | | | | | | ***10 s vs. 2 s*** | | | |
| | | | ***2 s*** | 0.095 | 0.006 | | | | | | | ***5 s vs. 2 s*** | | | |
| | | ***D5*** | ***10 s*** | 0.087 | 0.003 | Kruskal–Wallis H test | 3.69 | 2 | 0.158 | | Dunn's Test | ***10 s vs. 5 s*** | | | |
| | | | ***5 s*** | 0.100 | 0.006 | | | | | | | ***10 s vs. 2 s*** | | | |
| | | | ***2 s*** | 0.100 | 0.005 | | | | | | | ***5 s vs. 2 s*** | | | |
| | | ***D6*** | ***10 s*** | 0.101 | 0.006 | Kruskal–Wallis H test | 11.806 | 2 | 0.003 | ** | Dunn's Test | ***10 s vs. 5 s*** | 3.250 | 1.000 | |
| | | | ***5 s*** | 0.090 | 0.006 | | | | | | | ***10 s vs. 2 s*** | -11.750 | 0.003 | ** |
| | | | ***2 s*** | 0.122 | 0.003 | | | | | | | ***5 s vs. 2 s*** | -8.500 | 0.048 | * |
| | | ***D7*** | ***10 s*** | 0.092 | 0.005 | Kruskal–Wallis H test | 14.023 | 2 | 0.0009 | *** | Dunn's Test | ***10 s vs. 5 s*** | 4.438 | 0.626 | |
| | | | ***5 s*** | 0.082 | 0.003 | | | | | | | ***10 s vs. 2 s*** | -13.000 | 0.0007 | *** |
| | | | ***2 s*** | 0.113 | 0.004 | | | | | | | ***5 s vs. 2 s*** | -8.563 | 0.046 | * |
| | | ***D8*** | ***10 s*** | 0.079 | 0.005 | Kruskal–Wallis H test | 14.954 | 2 | 0.0006 | *** | Dunn's Test | ***10 s vs. 5 s*** | -0.375 | 1.000 | |
| | | | ***5 s*** | 0.082 | 0.006 | | | | | | | ***10 s vs. 2 s*** | -12.000 | 0.002 | ** |
| | | | ***2 s*** | 0.121 | 0.004 | | | | | | | ***5 s vs. 2 s*** | -11.625 | 0.003 | ** |
| | | ***D9*** | ***10 s*** | 0.053 | 0.006 | Kruskal–Wallis H test | 14.59 | 2 | 0.0007 | *** | Dunn's Test | ***10 s vs. 5 s*** | -10.063 | 0.013 | * |
| | | | ***5 s*** | 0.092 | 0.006 | | | | | | | ***10 s vs. 2 s*** | -12.813 | 0.0009 | *** |
| | | | ***2 s*** | 0.100 | 0.004 | | | | | | | ***5 s vs. 2 s*** | -2.750 | 1.000 | |
| | | ***D10*** | ***10 s*** | 0.071 | 0.008 | Kruskal–Wallis H test | 6.156 | 2 | 0.046 | * | Dunn's Test | ***10 s vs. 5 s*** | -4.000 | 0.772 | |
| | | | ***5 s*** | 0.084 | 0.008 | | | | | | | ***10 s vs. 2 s*** | -8.750 | 0.040 | * |
| | | | ***2 s*** | 0.100 | 0.004 | | | | | | | ***5 s vs. 2 s*** | -4.750 | 0.536 | |
| | | ***D11*** | ***10 s*** | 0.082 | 0.007 | Kruskal–Wallis H test | 5.959 | 2 | 0.051 | | Dunn's Test | ***10 s vs. 5 s*** | | | |
| | | | ***5 s*** | 0.098 | 0.007 | | | | | | | ***10 s vs. 2 s*** | | | |
| | | | ***2 s*** | 0.108 | 0.004 | | | | | | | ***5 s vs. 2 s*** | | | |
| | | ***D12*** | ***10 s*** | 0.069 | 0.006 | Kruskal–Wallis H test | 15.395 | 2 | 0.0005 | *** | Dunn's Test | ***10 s vs. 5 s*** | -2.375 | 1.000 | |
| | | | ***5 s*** | 0.073 | 0.005 | | | | | | | ***10 s vs. 2 s*** | -13.000 | 0.0007 | ** |
| | | | ***2 s*** | 0.118 | 0.004 | | | | | | | ***5 s vs. 2 s*** | -10.625 | 0.008 | *** |

*P < 0.05, **P < 0.01, and ***P < 0.001

| Table number | Related figure | Session | Group | Mean | SEM | Statistical test | Test Statistic | df | P | * | post-hoc test | Pair | Test Statistic | Bonferroni corrected P | * |
|---|---|---|---|---|---|---|---|---|---|---|---|---|---|---|---|
| Table S12 | Fig. S2A | D1 | 10 s | 2.88 | 1.22 | Kruskal–Wallis H test | 1.549 | 2 | 0.461 | | Dunn's Test | 10 s vs. 5 s | | | |
| | | | 5 s | 4.63 | 1.24 | | | | | | | 10 s vs. 2 s | | | |
| | | | 2 s | 3.75 | 0.77 | | | | | | | 5 s vs. 2 s | | | |
| | | D2 | 10 s | 1.50 | 0.85 | Kruskal–Wallis H test | 5.122 | 2 | 0.077 | | Dunn's Test | 10 s vs. 5 s | | | |
| | | | 5 s | 2.75 | 0.88 | | | | | | | 10 s vs. 2 s | | | |
| | | | 2 s | 3.50 | 0.57 | | | | | | | 5 s vs. 2 s | | | |
| | | D3 | 10 s | 2.38 | 0.71 | Kruskal–Wallis H test | 1.468 | 2 | 0.480 | | Dunn's Test | 10 s vs. 5 s | | | |
| | | | 5 s | 1.88 | 1.08 | | | | | | | 10 s vs. 2 s | | | |
| | | | 2 s | 1.38 | 0.26 | | | | | | | 5 s vs. 2 s | | | |
| | | D4 | 10 s | 2.75 | 0.77 | Kruskal–Wallis H test | 4.849 | 2 | 0.089 | | Dunn's Test | 10 s vs. 5 s | | | |
| | | | 5 s | 1.25 | 0.37 | | | | | | | 10 s vs. 2 s | | | |
| | | | 2 s | 1.00 | 0.19 | | | | | | | 5 s vs. 2 s | | | |
| | | D5 | 10 s | 4.88 | 0.99 | Kruskal–Wallis H test | 15.765 | 2 | 0.0004 | *** | Dunn's Test | 10 s vs. 5 s | -6.375 | 0.194 | |
| | | | 5 s | 2.13 | 0.55 | | | | | | | 10 s vs. 2 s | -13.688 | 0.0002 | *** |
| | | | 2 s | 0.50 | 0.19 | | | | | | | 5 s vs. 2 s | -7.313 | 0.102 | |
| | | D6 | 10 s | 2.75 | 0.70 | Kruskal–Wallis H test | 7.498 | 2 | 0.024 | * | Dunn's Test | 10 s vs. 5 s | -1.875 | 1.000 | |
| | | | 5 s | 2.13 | 0.44 | | | | | | | 10 s vs. 2 s | -8.625 | 0.028 | * |
| | | | 2 s | 1.00 | 0.19 | | | | | | | 5 s vs. 2 s | -6.750 | 0.125 | |
| | | D7 | 10 s | 3.50 | 0.63 | Kruskal–Wallis H test | 14.557 | 2 | 0.0007 | *** | Dunn's Test | 10 s vs. 5 s | -7.875 | 0.064 | |
| | | | 5 s | 1.63 | 0.65 | | | | | | | 10 s vs. 2 s | -12.938 | 0.0005 | *** |
| | | | 2 s | 0.50 | 0.19 | | | | | | | 5 s vs. 2 s | -5.063 | 0.416 | |
| | | D8 | 10 s | 4.63 | 1.08 | Kruskal–Wallis H test | 12.069 | 2 | 0.002 | ** | Dunn's Test | 10 s vs. 5 s | -7.500 | 0.069 | |
| | | | 5 s | 1.13 | 0.23 | | | | | | | 10 s vs. 2 s | -11.250 | 0.002 | ** |
| | | | 2 s | 0.63 | 0.18 | | | | | | | 5 s vs. 2 s | -3.750 | 0.766 | |
| | | D9 | 10 s | 4.63 | 0.75 | Kruskal–Wallis H test | 16.145 | 2 | 0.0003 | *** | Dunn's Test | 10 s vs. 5 s | -9.000 | 0.026 | * |
| | | | 5 s | 1.63 | 0.26 | | | | | | | 10 s vs. 2 s | -13.500 | 0.0002 | *** |
| | | | 2 s | 0.88 | 0.23 | | | | | | | 5 s vs. 2 s | -4.500 | 0.565 | |
| | | D10 | 10 s | 4.50 | 0.80 | Kruskal–Wallis H test | 18.289 | 2 | 0.0001 | *** | Dunn's Test | 10 s vs. 5 s | -10.000 | 0.007 | ** |
| | | | 5 s | 1.25 | 0.16 | | | | | | | 10 s vs. 2 s | -13.625 | 0.0001 | *** |
| | | | 2 s | 0.75 | 0.16 | | | | | | | 5 s vs. 2 s | -3.625 | 0.816 | |
| | | D11 | 10 s | 3.88 | 0.74 | Kruskal–Wallis H test | 16.653 | 2 | 0.0002 | *** | Dunn's Test | 10 s vs. 5 s | -7.250 | 0.095 | |
| | | | 5 s | 1.50 | 0.19 | | | | | | | 10 s vs. 2 s | -13.750 | 0.0001 | *** |
| | | | 2 s | 0.63 | 0.18 | | | | | | | 5 s vs. 2 s | -6.500 | 0.162 | |
| | | D12 | 10 s | 3.50 | 0.91 | Kruskal–Wallis H test | 12.427 | 2 | 0.002 | ** | Dunn's Test | 10 s vs. 5 s | -6.250 | 0.159 | |
| | | | 5 s | 1.25 | 0.16 | | | | | | | 10 s vs. 2 s | -11.375 | 0.001 | ** |
| | | | 2 s | 0.63 | 0.18 | | | | | | | 5 s vs. 2 s | -5.125 | 0.338 | |

*$P < 0.05$, **$P < 0.01$, and ***$P < 0.001$

| Table number | Related figure | Session | Group | Mean | SEM | Statistical test | Test Statistic | df | P | * | post-hoc test | Pair | Test Statistic | Bonferroni corrected P | * |
|---|---|---|---|---|---|---|---|---|---|---|---|---|---|---|---|
| Table S13 | Fig. S2A | D1 | 10 s | 0.48 | 0.29 | Kruskal–Wallis H test | 6.566 | 2 | 0.038 | * | Dunn's Test | 10 s vs. 5 s | -1.063 | 1.000 | |
| | | | 5 s | 0.56 | 0.23 | | | | | | | 10 s vs. 2 s | -8.313 | 0.056 | |
| | | | 2 s | 1.56 | 0.31 | | | | | | | 5 s vs. 2 s | -7.250 | 0.120 | |
| | | D2 | 10 s | 0.79 | 0.27 | Kruskal–Wallis H test | 7.361 | 2 | 0.025 | * | Dunn's Test | 10 s vs. 5 s | -3.625 | 0.915 | |
| | | | 5 s | 1.30 | 0.32 | | | | | | | 10 s vs. 2 s | -9.500 | 0.022 | * |
| | | | 2 s | 2.58 | 0.44 | | | | | | | 5 s vs. 2 s | -5.875 | 0.289 | |
| | | D3 | 10 s | 1.59 | 0.18 | Kruskal–Wallis H test | 11.37 | 2 | 0.003 | ** | Dunn's Test | 10 s vs. 5 s | -0.375 | 1.000 | |
| | | | 5 s | 1.57 | 0.36 | | | | | | | 10 s vs. 2 s | -10.500 | 0.009 | ** |
| | | | 2 s | 4.47 | 0.85 | | | | | | | 5 s vs. 2 s | -10.125 | 0.012 | * |
| | | D4 | 10 s | 2.21 | 0.39 | Kruskal–Wallis H test | 8.994 | 2 | 0.011 | * | Dunn's Test | 10 s vs. 5 s | -3.625 | 0.915 | |
| | | | 5 s | 2.96 | 0.50 | | | | | | | 10 s vs. 2 s | -10.438 | 0.009 | ** |
| | | | 2 s | 4.35 | 0.37 | | | | | | | 5 s vs. 2 s | -6.813 | 0.162 | |
| | | D5 | 10 s | 3.49 | 0.71 | Kruskal–Wallis H test | 1.165 | 2 | 0.559 | | Dunn's Test | 10 s vs. 5 s | | | |
| | | | 5 s | 3.28 | 0.66 | | | | | | | 10 s vs. 2 s | | | |
| | | | 2 s | 3.85 | 0.37 | | | | | | | 5 s vs. 2 s | | | |
| | | D6 | 10 s | 3.92 | 0.83 | Kruskal–Wallis H test | 0.635 | 2 | 0.728 | | Dunn's Test | 10 s vs. 5 s | | | |
| | | | 5 s | 4.38 | 0.66 | | | | | | | 10 s vs. 2 s | | | |
| | | | 2 s | 4.66 | 0.34 | | | | | | | 5 s vs. 2 s | | | |
| | | D7 | 10 s | 4.50 | 0.73 | Kruskal–Wallis H test | 0.903 | 2 | 0.637 | | Dunn's Test | 10 s vs. 5 s | | | |
| | | | 5 s | 4.31 | 0.30 | | | | | | | 10 s vs. 2 s | | | |
| | | | 2 s | 4.93 | 0.57 | | | | | | | 5 s vs. 2 s | | | |
| | | D8 | 10 s | 5.08 | 0.83 | Kruskal–Wallis H test | 0.466 | 2 | 0.792 | | Dunn's Test | 10 s vs. 5 s | | | |
| | | | 5 s | 4.80 | 0.68 | | | | | | | 10 s vs. 2 s | | | |
| | | | 2 s | 4.34 | 0.66 | | | | | | | 5 s vs. 2 s | | | |
| | | D9 | 10 s | 5.77 | 0.90 | Kruskal–Wallis H test | 0.815 | 2 | 0.665 | | Dunn's Test | 10 s vs. 5 s | | | |
| | | | 5 s | 4.74 | 0.53 | | | | | | | 10 s vs. 2 s | | | |
| | | | 2 s | 5.42 | 0.74 | | | | | | | 5 s vs. 2 s | | | |
| | | D10 | 10 s | 7.56 | 1.13 | Kruskal–Wallis H test | 3.518 | 2 | 0.172 | | Dunn's Test | 10 s vs. 5 s | | | |
| | | | 5 s | 5.56 | 0.19 | | | | | | | 10 s vs. 2 s | | | |
| | | | 2 s | 5.16 | 0.54 | | | | | | | 5 s vs. 2 s | | | |
| | | D11 | 10 s | 6.33 | 0.83 | Kruskal–Wallis H test | 1.24 | 2 | 0.538 | | Dunn's Test | 10 s vs. 5 s | | | |
| | | | 5 s | 4.97 | 0.61 | | | | | | | 10 s vs. 2 s | | | |
| | | | 2 s | 5.22 | 0.55 | | | | | | | 5 s vs. 2 s | | | |
| | | D12 | 10 s | 5.75 | 0.63 | Kruskal–Wallis H test | 0.035 | 2 | 0.983 | | Dunn's Test | 10 s vs. 5 s | | | |
| | | | 5 s | 5.91 | 0.47 | | | | | | | 10 s vs. 2 s | | | |
| | | | 2 s | 5.87 | 0.47 | | | | | | | 5 s vs. 2 s | | | |

*P < 0.05, **P < 0.01, and ***P < 0.001

| Table number | Related figure | Session | Group | Mean | SEM | Statistical test | Test Statistic | df | P | * | post-hoc test | Pair | Test Statistic | Bonferroni corrected P | * |
|---|---|---|---|---|---|---|---|---|---|---|---|---|---|---|---|
| ***Table S14*** | ***Fig. S2B*** | ***D1*** | ***10 s*** | -0.04 | 0.01 | Kruskal–Wallis H test | 5.218 | 2 | 0.074 | | Dunn's Test | ***10 s vs. 5 s*** | | | |
| | | | ***5 s*** | -0.07 | 0.01 | | | | | | | ***10 s vs. 2 s*** | | | |
| | | | ***2 s*** | -0.06 | 0.01 | | | | | | | ***5 s vs. 2 s*** | | | |
| | | ***D2*** | ***10 s*** | -0.04 | 0.01 | Kruskal–Wallis H test | 2.617 | 2 | 0.270 | | Dunn's Test | ***10 s vs. 5 s*** | | | |
| | | | ***5 s*** | 0.00 | 0.02 | | | | | | | ***10 s vs. 2 s*** | | | |
| | | | ***2 s*** | -0.03 | 0.02 | | | | | | | ***5 s vs. 2 s*** | | | |
| | | ***D3*** | ***10 s*** | -0.02 | 0.02 | Kruskal–Wallis H test | 10.151 | 2 | 0.006 | ** | Dunn's Test | ***10 s vs. 5 s*** | -2.563 | 1.000 | |
| | | | ***5 s*** | 0.01 | 0.02 | | | | | | | ***10 s vs. 2 s*** | -10.750 | 0.007 | ** |
| | | | ***2 s*** | 0.16 | 0.06 | | | | | | | ***5 s vs. 2 s*** | -8.188 | 0.061 | |
| | | ***D4*** | ***10 s*** | 0.03 | 0.02 | Kruskal–Wallis H test | 13.172 | 2 | 0.001 | ** | Dunn's Test | ***10 s vs. 5 s*** | -2.500 | 1.000 | |
| | | | ***5 s*** | 0.06 | 0.02 | | | | | | | ***10 s vs. 2 s*** | -12.125 | 0.002 | ** |
| | | | ***2 s*** | 0.24 | 0.03 | | | | | | | ***5 s vs. 2 s*** | -9.625 | 0.019 | * |
| | | ***D5*** | ***10 s*** | 0.11 | 0.05 | Kruskal–Wallis H test | 9.223 | 2 | 0.010 | * | Dunn's Test | ***10 s vs. 5 s*** | -0.188 | 1.000 | |
| | | | ***5 s*** | 0.09 | 0.03 | | | | | | | ***10 s vs. 2 s*** | -9.375 | 0.024 | * |
| | | | ***2 s*** | 0.26 | 0.03 | | | | | | | ***5 s vs. 2 s*** | -9.188 | 0.028 | * |
| | | ***D6*** | ***10 s*** | 0.13 | 0.06 | Kruskal–Wallis H test | 4.637 | 2 | 0.098 | | Dunn's Test | ***10 s vs. 5 s*** | | | |
| | | | ***5 s*** | 0.13 | 0.03 | | | | | | | ***10 s vs. 2 s*** | | | |
| | | | ***2 s*** | 0.27 | 0.04 | | | | | | | ***5 s vs. 2 s*** | | | |
| | | ***D7*** | ***10 s*** | 0.13 | 0.06 | Kruskal–Wallis H test | 7.949 | 2 | 0.019 | * | Dunn's Test | ***10 s vs. 5 s*** | -1.438 | 1.000 | |
| | | | ***5 s*** | 0.15 | 0.03 | | | | | | | ***10 s vs. 2 s*** | -9.250 | 0.026 | * |
| | | | ***2 s*** | 0.30 | 0.02 | | | | | | | ***5 s vs. 2 s*** | -7.813 | 0.081 | |
| | | ***D8*** | ***10 s*** | 0.18 | 0.06 | Kruskal–Wallis H test | 3.502 | 2 | 0.174 | | Dunn's Test | ***10 s vs. 5 s*** | | | |
| | | | ***5 s*** | 0.18 | 0.05 | | | | | | | ***10 s vs. 2 s*** | | | |
| | | | ***2 s*** | 0.27 | 0.03 | | | | | | | ***5 s vs. 2 s*** | | | |
| | | ***D9*** | ***10 s*** | 0.29 | 0.07 | Kruskal–Wallis H test | 2.641 | 2 | 0.267 | | Dunn's Test | ***10 s vs. 5 s*** | | | |
| | | | ***5 s*** | 0.17 | 0.05 | | | | | | | ***10 s vs. 2 s*** | | | |
| | | | ***2 s*** | 0.28 | 0.05 | | | | | | | ***5 s vs. 2 s*** | | | |
| | | ***D10*** | ***10 s*** | 0.33 | 0.08 | Kruskal–Wallis H test | 4.74 | 2 | 0.093 | | Dunn's Test | ***10 s vs. 5 s*** | | | |
| | | | ***5 s*** | 0.15 | 0.03 | | | | | | | ***10 s vs. 2 s*** | | | |
| | | | ***2 s*** | 0.30 | 0.04 | | | | | | | ***5 s vs. 2 s*** | | | |
| | | ***D11*** | ***10 s*** | 0.28 | 0.07 | Kruskal–Wallis H test | 7.408 | 2 | 0.025 | * | Dunn's Test | ***10 s vs. 5 s*** | -6.875 | 0.154 | |
| | | | ***5 s*** | 0.14 | 0.02 | | | | | | | ***10 s vs. 2 s*** | -2.375 | 1.000 | |
| | | | ***2 s*** | 0.32 | 0.02 | | | | | | | ***5 s vs. 2 s*** | -9.250 | 0.026 | * |
| | | ***D12*** | ***10 s*** | 0.26 | 0.06 | Kruskal–Wallis H test | 5.959 | 2 | 0.051 | | Dunn's Test | ***10 s vs. 5 s*** | | | |
| | | | ***5 s*** | 0.17 | 0.02 | | | | | | | ***10 s vs. 2 s*** | | | |
| | | | ***2 s*** | 0.33 | 0.02 | | | | | | | ***5 s vs. 2 s*** | | | |

**P < 0.05, **P < 0.01, and ***P < 0.001*

| Table number | Related figure | Session | Group | Mean | SEM | Statistical test | Test Statistic | df | P | * | post-hoc test | Pair | Test Statistic | Bonferroni corrected P | * |
|---|---|---|---|---|---|---|---|---|---|---|---|---|---|---|---|
| ***Table S15*** | ***Fig. 6C and S3A*** | ***D1*** | ***10 s*** | 0.11 | 0.05 | Kruskal–Wallis H test | 0.84 | 2 | 0.657 | | Dunn's Test | ***10 s vs. 5 s*** | | | |
| | | | ***5 s*** | 0.05 | 0.03 | | | | | | | ***10 s vs. 2 s*** | | | |
| | | | ***2 s*** | 0.04 | 0.02 | | | | | | | ***5 s vs. 2 s*** | | | |
| | | ***D2*** | ***10 s*** | 0.28 | 0.08 | Kruskal–Wallis H test | 3.698 | 2 | 0.157 | | Dunn's Test | ***10 s vs. 5 s*** | | | |
| | | | ***5 s*** | 0.38 | 0.09 | | | | | | | ***10 s vs. 2 s*** | | | |
| | | | ***2 s*** | 0.15 | 0.03 | | | | | | | ***5 s vs. 2 s*** | | | |
| | | ***D3*** | ***10 s*** | 0.90 | 0.15 | Kruskal–Wallis H test | 4.436 | 2 | 0.109 | | Dunn's Test | ***10 s vs. 5 s*** | | | |
| | | | ***5 s*** | 0.53 | 0.12 | | | | | | | ***10 s vs. 2 s*** | | | |
| | | | ***2 s*** | 0.50 | 0.11 | | | | | | | ***5 s vs. 2 s*** | | | |
| | | ***D4*** | ***10 s*** | 1.43 | 0.26 | Kruskal–Wallis H test | 7.656 | 2 | 0.022 | * | Dunn's Test | ***10 s vs. 5 s*** | -1.500 | 0.094 | |
| | | | ***5 s*** | 0.75 | 0.13 | | | | | | | ***10 s vs. 2 s*** | -9.000 | 0.029 | * |
| | | | ***2 s*** | 0.68 | 0.06 | | | | | | | ***5 s vs. 2 s*** | -7.500 | 1.000 | |
| | | ***D5*** | ***10 s*** | 1.93 | 0.45 | Kruskal–Wallis H test | 5.844 | 2 | 0.054 | | Dunn's Test | ***10 s vs. 5 s*** | | | |
| | | | ***5 s*** | 0.96 | 0.18 | | | | | | | ***10 s vs. 2 s*** | | | |
| | | | ***2 s*** | 0.73 | 0.05 | | | | | | | ***5 s vs. 2 s*** | | | |
| | | ***D6*** | ***10 s*** | 2.34 | 0.56 | Kruskal–Wallis H test | 5.879 | 2 | 0.053 | | Dunn's Test | ***10 s vs. 5 s*** | | | |
| | | | ***5 s*** | 1.08 | 0.17 | | | | | | | ***10 s vs. 2 s*** | | | |
| | | | ***2 s*** | 0.79 | 0.07 | | | | | | | ***5 s vs. 2 s*** | | | |
| | | ***D7*** | ***10 s*** | 2.18 | 0.56 | Kruskal–Wallis H test | 5.594 | 2 | 0.061 | | Dunn's Test | ***10 s vs. 5 s*** | | | |
| | | | ***5 s*** | 1.15 | 0.16 | | | | | | | ***10 s vs. 2 s*** | | | |
| | | | ***2 s*** | 0.83 | 0.05 | | | | | | | ***5 s vs. 2 s*** | | | |
| | | ***D8*** | ***10 s*** | 2.56 | 0.60 | Kruskal–Wallis H test | 12.009 | 2 | 0.002 | ** | Dunn's Test | ***10 s vs. 5 s*** | -6.000 | 0.236 | |
| | | | ***5 s*** | 1.29 | 0.26 | | | | | | | ***10 s vs. 2 s*** | -12.188 | 0.002 | ** |
| | | | ***2 s*** | 0.78 | 0.05 | | | | | | | ***5 s vs. 2 s*** | -6.188 | 0.264 | |
| | | ***D9*** | ***10 s*** | 3.39 | 0.76 | Kruskal–Wallis H test | 12.431 | 2 | 0.002 | ** | Dunn's Test | ***10 s vs. 5 s*** | -6.063 | 0.221 | |
| | | | ***5 s*** | 1.29 | 0.23 | | | | | | | ***10 s vs. 2 s*** | -12.313 | 0.001 | ** |
| | | | ***2 s*** | 0.75 | 0.09 | | | | | | | ***5 s vs. 2 s*** | -6.250 | 0.248 | |
| | | ***D10*** | ***10 s*** | 4.00 | 0.87 | Kruskal–Wallis H test | 10.581 | 2 | 0.005 | ** | Dunn's Test | ***10 s vs. 5 s*** | -5.250 | 0.236 | |
| | | | ***5 s*** | 1.18 | 0.16 | | | | | | | ***10 s vs. 2 s*** | -11.438 | 0.003 | ** |
| | | | ***2 s*** | 0.80 | 0.07 | | | | | | | ***5 s vs. 2 s*** | -6.188 | 0.408 | |
| | | ***D11*** | ***10 s*** | 3.59 | 0.71 | Kruskal–Wallis H test | 13.851 | 2 | 0.00098 | *** | Dunn's Test | ***10 s vs. 5 s*** | -6.813 | 0.225 | |
| | | | ***5 s*** | 1.20 | 0.10 | | | | | | | ***10 s vs. 2 s*** | -13.063 | 0.0006 | *** |
| | | | ***2 s*** | 0.85 | 0.03 | | | | | | | ***5 s vs. 2 s*** | -6.250 | 0.157 | |
| | | ***D12*** | ***10 s*** | 3.33 | 0.59 | Kruskal–Wallis H test | 17.786 | 2 | 0.0001 | *** | Dunn's Test | ***10 s vs. 5 s*** | -7.313 | 0.098 | |
| | | | ***5 s*** | 1.21 | 0.10 | | | | | | | ***10 s vs. 2 s*** | -14.813 | 0.0001 | *** |
| | | | ***2 s*** | 0.89 | 0.04 | | | | | | | ***5 s vs. 2 s*** | -7.500 | 0.112 | |

*$P < 0.05$, **$P < 0.01$, and ***$P < 0.001$

| Table number | Related figure | Session | Group | Component | Mean | Statistical test | Correlation coffeficient | P | * |
|---|---|---|---|---|---|---|---|---|---|
| Table S16 | Fig. 6A, 6B, and S3B | D1 | 10 s | Number of HE* | 11.25 | Spearman test | 1 | 0 | *** |
| | | | | HE accuracy | 8.75 | | | | |
| | | | 5 s | Number of HE* | 5.00 | Spearman test | 1.000 | 0.0001 | *** |
| | | | | HE accuracy | 5.00 | | | | |
| | | | 2 s | Number of HE* | 3.75 | Spearman test | 1 | 0 | *** |
| | | | | HE accuracy | 3.75 | | | | |
| | | D2 | 10 s | Number of HE* | 27.50 | Spearman test | 0.975 | 0.00004 | *** |
| | | | | HE accuracy | 22.50 | | | | |
| | | | 5 s | Number of HE* | 37.50 | Spearman test | 0.963 | 0.0001 | *** |
| | | | | HE accuracy | 36.25 | | | | |
| | | | 2 s | Number of HE* | 15.00 | Spearman test | 0.900 | 0.002 | ** |
| | | | | HE accuracy | 13.75 | | | | |
| | | D3 | 10 s | Number of HE* | 90.00 | Spearman test | 0.877 | 0.004 | ** |
| | | | | HE accuracy | 61.25 | | | | |
| | | | 5 s | Number of HE* | 52.50 | Spearman test | 0.946 | 0.0004 | *** |
| | | | | HE accuracy | 47.50 | | | | |
| | | | 2 s | Number of HE* | 50.00 | Spearman test | 1 | 0 | *** |
| | | | | HE accuracy | 42.50 | | | | |
| | | D4 | 10 s | Number of HE* | 142.50 | Spearman test | 0.786 | 0.021 | * |
| | | | | HE accuracy | 81.25 | | | | |
| | | | 5 s | Number of HE* | 75.00 | Spearman test | 0.938 | 0.0006 | *** |
| | | | | HE accuracy | 63.75 | | | | |
| | | | 2 s | Number of HE* | 67.50 | Spearman test | 0.850 | 0.007 | ** |
| | | | | HE accuracy | 60.00 | | | | |
| | | D5 | 10 s | Number of HE* | 192.50 | Spearman test | 0.661 | 0.074 | |
| | | | | HE accuracy | 85.00 | | | | |
| | | | 5 s | Number of HE* | 96.25 | Spearman test | 0.994 | 0.000001 | *** |
| | | | | HE accuracy | 77.50 | | | | |
| | | | 2 s | Number of HE* | 72.50 | Spearman test | 0.882 | 0.004 | ** |
| | | | | HE accuracy | 68.75 | | | | |
| | | D6 | 10 s | Number of HE* | 233.75 | Spearman test | 0.768 | 0.026 | * |
| | | | | HE accuracy | 91.25 | | | | |
| | | | 5 s | Number of HE* | 107.50 | Spearman test | 0.870 | 0.005 | ** |
| | | | | HE accuracy | 82.50 | | | | |
| | | | 2 s | Number of HE* | 78.75 | Spearman test | 0.894 | 0.003 | ** |
| | | | | HE accuracy | 72.50 | | | | |
| | | D7 | 10 s | Number of HE* | 217.50 | Spearman test | 0.619 | 0.102 | |
| | | | | HE accuracy | 88.75 | | | | |
| | | | 5 s | Number of HE* | 115.00 | Spearman test | 0.474 | 0.235 | |
| | | | | HE accuracy | 90.00 | | | | |
| | | | 2 s | Number of HE* | 82.50 | Spearman test | 0.384 | 0.348 | |
| | | | | HE accuracy | 72.50 | | | | |
| | | D8 | 10 s | Number of HE* | 256.25 | Spearman test | 0.378 | 0.356 | |
| | | | | HE accuracy | 97.50 | | | | |
| | | | 5 s | Number of HE* | 128.75 | Spearman test | 0.739 | 0.036 | * |
| | | | | HE accuracy | 88.75 | | | | |
| | | | 2 s | Number of HE* | 77.50 | Spearman test | 0.963 | 0.0001 | *** |
| | | | | HE accuracy | 76.25 | | | | |
| | | D9 | 10 s | Number of HE* | 338.75 | Spearman test | 0.646 | 0.083 | |
| | | | | HE accuracy | 93.75 | | | | |
| | | | 5 s | Number of HE* | 128.75 | Spearman test | 0.809 | 0.015 | * |
| | | | | HE accuracy | 92.50 | | | | |
| | | | 2 s | Number of HE* | 75.00 | Spearman test | 0.890 | 0.003 | ** |
| | | | | HE accuracy | 71.25 | | | | |
| | | D10 | 10 s | Number of HE* | 400.00 | Spearman test | 0.866 | 0.005 | ** |
| | | | | HE accuracy | 93.75 | | | | |
| | | | 5 s | Number of HE* | 117.50 | Spearman test | 0.803 | 0.016 | * |
| | | | | HE accuracy | 91.25 | | | | |
| | | | 2 s | Number of HE* | 80.00 | Spearman test | 0.969 | 0.00007 | *** |
| | | | | HE accuracy | 76.25 | | | | |
| | | D11 | 10 s | Number of HE* | 358.75 | Spearman test | 0.845 | 0.008 | ** |
| | | | | HE accuracy | 96.25 | | | | |
| | | | 5 s | Number of HE* | 120.00 | Spearman test | 0.548 | 0.160 | |
| | | | | HE accuracy | 93.75 | | | | |
| | | | 2 s | Number of HE* | 85.00 | Spearman test | 0.960 | 0.00016 | *** |
| | | | | HE accuracy | 83.75 | | | | |
| | | D12 | 10 s | Number of HE* | 332.50 | Spearman test | -0.113 | 0.789 | |
| | | | | HE accuracy | 96.25 | | | | |
| | | | 5 s | Number of HE* | 121.25 | Spearman test | 0.394 | 0.334 | |
| | | | | HE accuracy | 95.00 | | | | |
| | | | 2 s | Number of HE* | 88.75 | Spearman test | 0.855 | 0.007 | ** |
| | | | | HE accuracy | 86.25 | | | | |

\ Number of HE* = Number of HE * 100 *P < 0.05, **P < 0.01, and ***P < 0.001
